# A genome-wide atlas of human cell morphology

**DOI:** 10.1101/2023.08.06.552164

**Authors:** Meraj Ramezani, Julia Bauman, Avtar Singh, Erin Weisbart, John Yong, Maria Lozada, Gregory P. Way, Sanam L. Kavari, Celeste Diaz, Marzieh Haghighi, Thiago M. Batista, Joaquín Pérez-Schindler, Melina Claussnitzer, Shantanu Singh, Beth A. Cimini, Paul C. Blainey, Anne E. Carpenter, Calvin H. Jan, James T. Neal

## Abstract

A key challenge of the modern genomics era is developing data-driven representations of gene function. Here, we present the first unbiased morphology-based genome-wide perturbation atlas in human cells, containing three genome-scale genotype-phenotype maps comprising >20,000 single-gene CRISPR-Cas9-based knockout experiments in >30 million cells. Our optical pooled cell profiling approach (PERISCOPE) combines a de-stainable high-dimensional phenotyping panel (based on Cell Painting^1,2^) with optical sequencing of molecular barcodes and a scalable open-source analysis pipeline to facilitate massively parallel screening of pooled perturbation libraries. This approach provides high-dimensional phenotypic profiles of individual cells, while simultaneously enabling interrogation of subcellular processes. Our atlas reconstructs known pathways and protein-protein interaction networks, identifies culture media-specific responses to gene knockout, and clusters thousands of human genes by phenotypic similarity. Using this atlas, we identify the poorly-characterized disease-associated transmembrane protein TMEM251/LYSET as a Golgi-resident protein essential for mannose-6-phosphate-dependent trafficking of lysosomal enzymes, showing the power of these representations. In sum, our atlas and screening technology represent a rich and accessible resource for connecting genes to cellular functions at scale.

## Introduction

Large scale DNA sequencing has transformed our ability to identify and catalog diverse genotypic information but created a new bottleneck: characterizing the diverse impacts of genotype on human biology. Thus, systematically connecting human genes and genotypes to disease- and trait-relevant phenotypes remains a grand challenge for biomedicine.

Pooled CRISPR screens^3^ have proven a powerful tool for tackling this challenge, but typically require compromising on either phenotypic content or scale. Genome-scale pooled CRISPR screens enable systematic assessment of gene function but compatible phenotypes, such as proliferation or cell death, are often simple or require a targeted assay, making them inappropriate for assessing many biologically-relevant processes in human cells, which are often subtle, graded, and/or complex^4^. In contrast, high-content profiling approaches such as imaging, transcriptomics, proteomics, and metabolomics can capture hundreds of quantitative phenotypes for each sample, providing a rich phenotypic profile, but are typically incompatible with genome-scale perturbation. A notable exception is Perturb-seq^5–8^, which has very recently been applied to profile the effects of CRISPRi knockdown of the expressed genome of the human chronic myeloid leukemia cell line K562^9^. This study demonstrated the immense value of generating rich, high-dimensional representations of cell state at genome scale using a new (and not yet widely available) DNA sequencing technology^10,11^ and resource-intensive data generation effort.

Optical pooled screening, which combines image-based phenotyping with image-based sequencing of perturbation barcodes, has emerged as a promising and complementary approach for high-dimensional genotype-phenotype mapping at single cell resolution that is scalable and cost-effective^12–15^. Optical pooled screens enable quantitative assessment of phenotypes invisible to molecular profiling approaches, such as cell morphology and subcellular localization, with greater throughput than arrayed image-based screens^16^, and, in contrast to pooled enrichment-based imaging approaches^17–20^, have no requirement for physical selection or predefinition of phenotypes.

Here, we combined high-dimensional multiparameter image-based cell phenotyping with massively-parallel optical pooled CRISPR screens in order to build the first genome-scale perturbation atlas of morphology phenotypes in human cells. We report the design of an optimized cell phenotyping panel based on the popular Cell Painting ^1,2^ image-based profiling assay that enables 5-color fluorescence microscopy of cell phenotypes followed by 4-color *in situ* sequencing-by-synthesis (ISS) to assign perturbations to cells. We also built a scalable, open-source, cloud-based pipeline for generating barcoded image-based profiles from genome-scale perturbation datasets. We use this technology to execute a whole-genome pooled optical CRISPR screen in human lung cancer cells (A549), profiling the effects of >20,000 single gene knockout experiments in unbiased fashion. Finally, we apply our approach to map genome-wide gene-by-environment interactions by executing two additional whole genome screens in human cervical cancer cells (HeLa) cultured either in traditional cell culture media or physiologic media^21^. Together, this work establishes a comprehensive resource for connecting human genotypes to high-dimensional image-based cellular phenotypes at scale.

## Results

### A genome-scale approach for high-dimensional optical CRISPR screens

To assess genome-wide knockout effects on cell morphology, we first constructed a whole genome CRISPR guide RNA library optimized for optical screening. To build this library, we selected on average four sgRNAs per gene from existing libraries^22,23^, identifying sgRNA sequences that would allow for total deconvolution of the sgRNA library in 12 cycles of *in situ* sequencing while also allowing for a Levenshtein distance of 2 between sgRNA sequences to enable error detection^24^, resulting in a library containing 80,862 sgRNAs targeting 20,393 genes (Supplementary Table 1). We cloned the sgRNA library into the CROP-seq vector^7^, enabling expression and direct *in situ* sequencing of sgRNA sequences (interchangeably referred to as barcodes), and packaged it for lentiviral delivery.

To comprehensively map genome-wide gene knockout effects to high-dimensional image-based phenotypes, we built a high-throughput data generation and analysis pipeline, PERISCOPE (Perturbation Effect Readout In situ with Single Cell Optical Phenotyping), comprising a suite of highly scalable wet and dry lab protocols that enables facile screening of genome-scale perturbation libraries by optical profiling. We first developed an optimized, de-stainable variant of the Cell Painting panel to collect morphological data by fluorescence imaging of cell compartments, followed by *in situ* sequencing of sgRNAs to assign perturbations to cells (Fig. 1a). This approach results in five phenotypic images for each cell – Phalloidin (Actin), anti-TOMM20 antibody (mitochondria), WGA (Golgi and cell membrane), ConA (Endoplasmic Reticulum) and DAPI (Nucleus) – plus 12 sequencing images, which are used to identify sequential sgRNA bases (Fig. 1b + Extended Data Fig. 1). To overcome spectral overlap between fluorescent phenotyping markers and fluorescent sequencing signal, we conjugated phenotypic probes to fluorophores using a disulfide linker^25,26^. This strategy allows five-color labeling followed by treatment with TCEP, a reducing agent, resulting in linker cleavage and liberation of linked fluorophores, freeing up fluorescent channels for *in situ* sequencing (Fig. 1c). To analyze these data, we modified the standard Cell Painting image analysis workflow within the open source image analysis software CellProfiler^27^ to handle the added complexity of pooled perturbations, including the incorporation of image alignment across different resolutions and barcode calling^14^ (Fig. 1d). Similarly, we adapted our data analysis workflow based on the open-source PyCytominer library (see Methods) to process single-cell profiles using pooled data rather than arrayed data.

**Figure 1.**
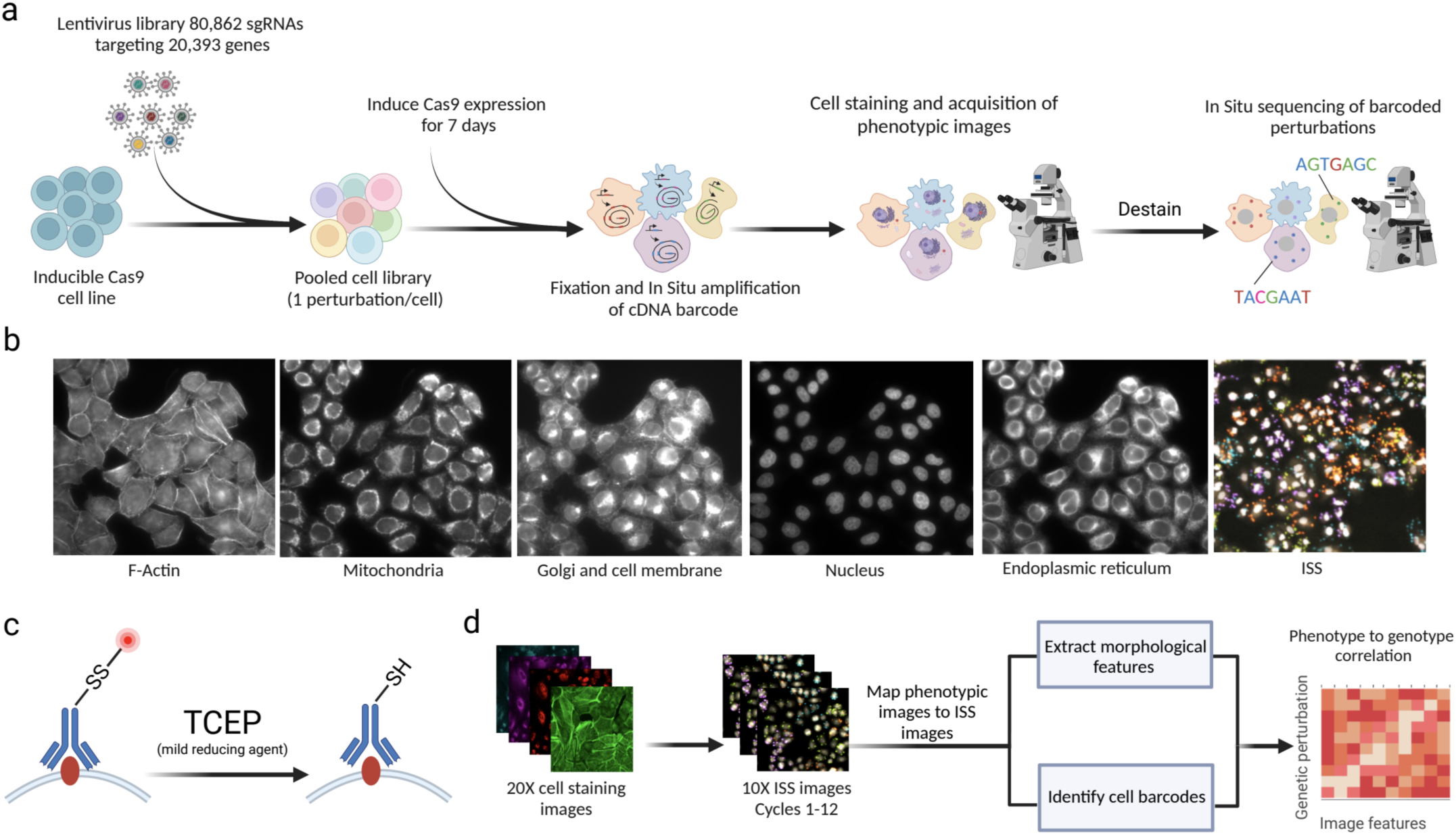
Pooled optical screens with PERISCOPE. (a) Experimental workflow for PERISCOPE screens. (b) Example images of five phenotypic stains and fluorescent in situ sequencing. (c) Schematic of destaining strategy to enable in situ sequencing after fluorescence imaging of phenotypic stains.(d) Overview of the PERISCOPE analysis pipeline including extraction of phenotypic features, deconvolution of barcodes and genotype-phenotype correlation. Schematics created with Biorender.

### A morphology-based genome-wide perturbation map in human lung cancer cells

We first aimed to demonstrate the scalability and robustness of the PERISCOPE pipeline by executing a whole genome pooled optical CRISPR screen in human lung cancer cells (A549). Using 48 identically-prepared wells of 6-well plates, we collected morphological profiles from 11,211,357 single cells which yielded 20,393 gene-level profiles at an average representation of 460 cells per gene (Extended Data Fig. 2a). Crucially, optical sgRNA counts were highly correlated with counts obtained from next-generation sequencing of perturbed cells (Extended Data Fig. 2b), validating *in situ* sequencing accuracy, and between biological screen replicates, validating screen robustness (Extended Data Fig. 2c-e). As expected, the PERISCOPE pipeline reported that perturbation of TOMM20, the direct target of the antibody stain for mitochondria, impacted the expected mitochondrial features (Extended Data Fig. 2f).

We next applied a hit calling pipeline that we designed to identify gene perturbation signatures above background noise using image-based features. Optical profiling collects spatial information and thus our pipeline was able to identify two classes of screen hit – “whole cell” hit genes, which were defined based upon aggregate signal from all cell compartments, and “compartment” hit genes identified by imaging measurements from a subset of the five labeled subcellular compartments (see Methods). Overall, we identified 1,339 whole-cell hit genes, and 2,320 compartment hit genes, for a total of 3,659 hits (Fig. 2a, Supplementary Table 2). We found roughly equal numbers of compartment hit genes in each subcellular compartment, demonstrating that each channel is providing useful information (Fig. 2b and Extended Data Fig. 3e-f). Importantly, we also observed that knocking out genes known to act in well-defined cell compartment-specific roles produced strong morphological phenotypes in those compartments. Specifically, we selected genes encoding five compartment-associated protein complex members and grouped their morphological profiles by complex. For each of these complexes, we observed an enrichment in phenotypic features extracted from the expected cellular compartment (Fig. 2c). For example, while perturbations targeting outer mitochondrial membrane (OMM) proteins produce morphological phenotypes throughout the cell, a plurality (32%) of the overall signal is concentrated in the mitochondria. Likewise, sgRNAs targeting genes involved in protein mannosylation display an enrichment in phenotypic features from the endoplasmic reticulum (ER), where synthesis of mannosyl donor substrates and mannosyltransfer to proteins takes place^28^.

**Figure 2.**
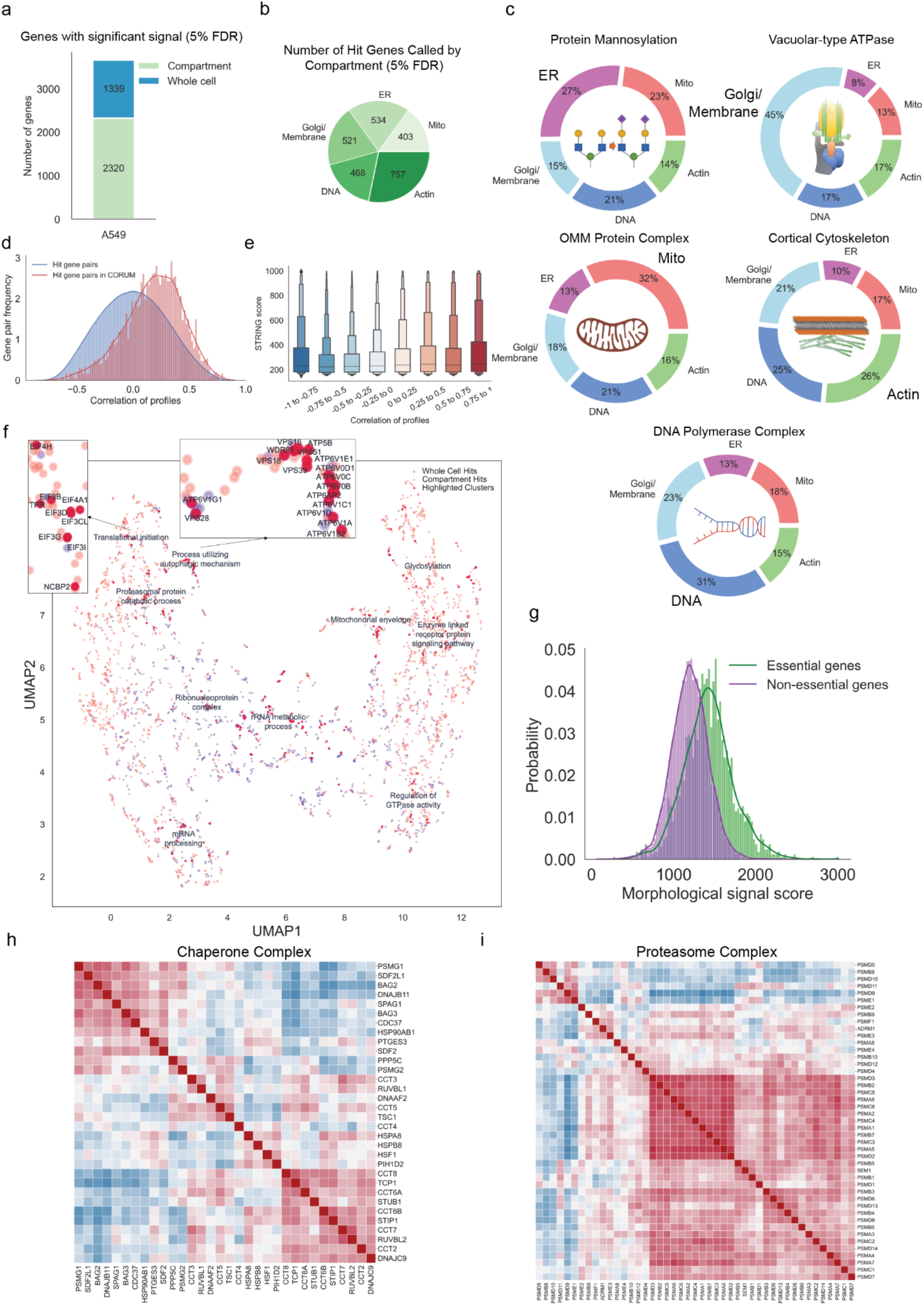
A genome-wide perturbation map in A549 cells. Summary of whole genome PERISCOPE screen performed in A549 cells. (a) Hit genes identified in the screen include some single-compartment and some impacting multiple compartments and features across the cell. Green represents hit genes called based on a subset of cell compartments (ER, mitochondria, actin, DNA and Golgi/membrane) and blue represents hit genes called based on overall gene profile. Detailed description in the methods section. (b) Hit genes called based on a single compartment are distributed across all five measured compartments. It is possible for a gene to be hit in multiple compartments without being a whole cell hit, see Extended Data Fig. 3a-b for more details. (c) Pie charts showing the average normalized fraction of number of features significantly different from the control categorized based on target compartments for genes in the indicated set.(d) Distributions of optical profile correlations among all possible gene pairs versus correlations among gene pairs representing CORUM4.0 protein complexes that have at least two thirds of complex subunits within hit genes. (e) Boxen (Letter-value) plot representing STRING scores divided into bins based on PERISCOPE profile correlation between gene pairs. (f) UMAP embedding of the hit gene profiles from the A549 dataset. Each dot represents a genetic perturbation and distance implies the correlation of profile in a two dimensional embedding. Manual annotation of cluster functions are presented for highlighted clusters based on gene ontology (GO) data sets. Example insets show coherent clustering of related genes. (g) The distribution of morphological signal scores for essential and nonessential genes (DepMap gene effect at -0.5 threshold) for all perturbations in the A549 dataset. (h, i) Heatmaps representing Pearson correlation between gene profiles after hierarchical clustering using Ward’s method. Gene complexes/processes were enriched in the A549 dataset based on the preranked GSEA analysis. (h) displays hit genes belonging to the GO:CC chaperone complex genes set (GO:0101031). (i) displays hit genes belonging to the GO:CC proteasome complex (GO:0000502).

We next benchmarked image-based gene knockout profiles against existing databases of gene function. First, using profile correlation between gene knockouts as a proxy for functional similarity between genes, we compared our screen data to the protein-protein interaction databases CORUM^29^ and STRING^30^. Of 3,659 total hits, we identified 1,271 genes belonging to 501 unique complexes present in the CORUM4.0 database. Profiles from hit gene pairs within a cluster showed higher correlation values than the background distribution of all possible hit gene pairs (Fig. 2d). Additionally, morphological profile pairs with higher correlations demonstrated higher protein-protein interaction confidence scores from the STRING database (Fig. 2e). We subsequently evaluated the extent to which image-based gene knockout profiles were correlated with gene knockout fitness effects using the Broad Institute’s Dependency Map (DepMap) database^31^. While essential genes were more likely on average to produce a high signal score (see Methods), the majority of screen hits (72%) were nonessential genes, consistent with most gene knockouts producing optical phenotypes beyond simple cell toxicity (Fig. 2g & Extended Data Fig. 4a). Further, morphological signal score was not well correlated with baseline gene expression, with many genes expressed at low levels still producing significant morphological signal when perturbed demonstrating orthogonality of optical phenotypes (Extended Data Fig 4d).

We performed unbiased clustering of screen hits based on morphological similarity and visualized high-level similarity between morphological profiles via 2-dimensional UMAP embedding (Fig 2.f). We observed logical clustering by biological function across an array of processes, such as translational initiation, lysosome acidification, autophagy, proteasomal protein catabolic processes, mRNA processing, rRNA metabolic process, glycosylation, regulation of GTPase activity and others. Hierarchical clustering based on high dimensional profiles also revealed biologically coherent clustering of perturbations targeting related genes. For example, we identified a cluster of molecular chaperones (Fig. 2h) displaying high similarity between sgRNAs targeting CCT genes that form the chaperonin-containing TCP1 complex, which is essential for producing native actin, tubulin, and other proteins involved in cell cycle progression.^32^

We also observed biologically coherent similarity within the family of genes encoding proteins that form the proteasome (Fig. 2i). The mammalian proteasome is a large protein degradation complex that exists in multiple configurations: the canonical proteasome consists of the 20S catalytic core particle capped on either end by the 19S regulatory particle and the interferon-inducible immunoproteasome, which replaces several catalytic subunits in the 20S and can be alternatively capped by the 11S particle^33^. We observed a highly correlated subcluster of 30 screen hits, representing mostly essential genes in the A549 cell line (average -1.63 Chronos gene effect score^34^). The alpha and beta subunits of the catalytic core particle, as well as the ATPase subunits of the regulatory 19S particle, showed the strongest levels of correlation in our data, whereas the non-ATPase subunits of the regulatory particle exhibited lower signal. A notable exception is PSMD14/Rpn11, which de-ubiquiylates substrates as they enter the proteasome and clustered with the catalytic core. Blocking de-ubiquitylation stalls substrate entry, thus impairing overall proteasome function, consistent with the morphological similarity observed for catalytic subunits responsible for substrate translocation and degradation ^35^. As expected, the interferon gamma-inducible subunits of both the catalytic core particle (PSMB8, PSMB9, and PSMB10) and 11S particle (PSME1, PSME2) displayed weak signal in the absence of interferon stimulus and did not correlate with core proteasome components.

### Mapping gene-by-environment interactions at genome scale

Cell metabolism is influenced by a vast array of interactions between genes and environmental stimuli, and as such, *in vitro* genetic screens carried out in traditional cell culture media, which poorly recapitulate physiologic environments, may fail to capture metabolically relevant phenotypes. Recently, “physiologic media” such as Plasmax^36^ or human plasma-like medium (HPLM)^37^ have been developed as tools to study the effects of genetic perturbations under environmental conditions designed to more accurately mimic in vivo human physiology, and in a recent study, HPLM was shown to dramatically alter the spectrum of gene essentiality in K562 cells^21^. Such studies demonstrate the usefulness of screening under physiologically relevant conditions, but have been limited to growth assays, preventing the systematic assessment of gene perturbation on high-dimensional cell phenotypes.

Having demonstrated the scalability and robustness of the PERISCOPE technique as well as its ability to capture biological relationships based on high-dimensional morphological profiles, we next applied PERISCOPE to investigate gene-by-environment interactions at genome scale. To accomplish this, we performed two genome-scale PERISCOPE screens (as described above) in HeLa cells cultured in either Dulbecco’s Modified Eagle Medium (DMEM) or HPLM. In addition to their experimental tractability and prior validation in optical screening workflows^12–14^, HeLa cells have been demonstrated to exhibit sensitivity to metabolic environmental cues such as altered glucose levels^38,39^. In our screens, we collected single-cell morphological profiles from 12,319,688 (mean 491 cells per gene) and 9,118,260 (366 cells per gene) individual cells in the DMEM and HPLM screens, respectively. We identified a similar number of hit genes in both types of media, representing ∼42% of the expressed genome (4,712 hit genes in the DMEM screen: 2074 common, 2638 unique and 4,771 in HPLM: 2697 unique) (Fig. 3a, Supplementary Table 2). We again identified compartment hits from all subcellular compartments (Fig. 3b, Extended Data Fig. 3a-d), and as in the A549 screen we found that physically interacting proteins (per CORUM and STRING) were more likely to have similar morphological profiles than random hit gene pairs (Fig. 3c,d).

**Figure 3.**
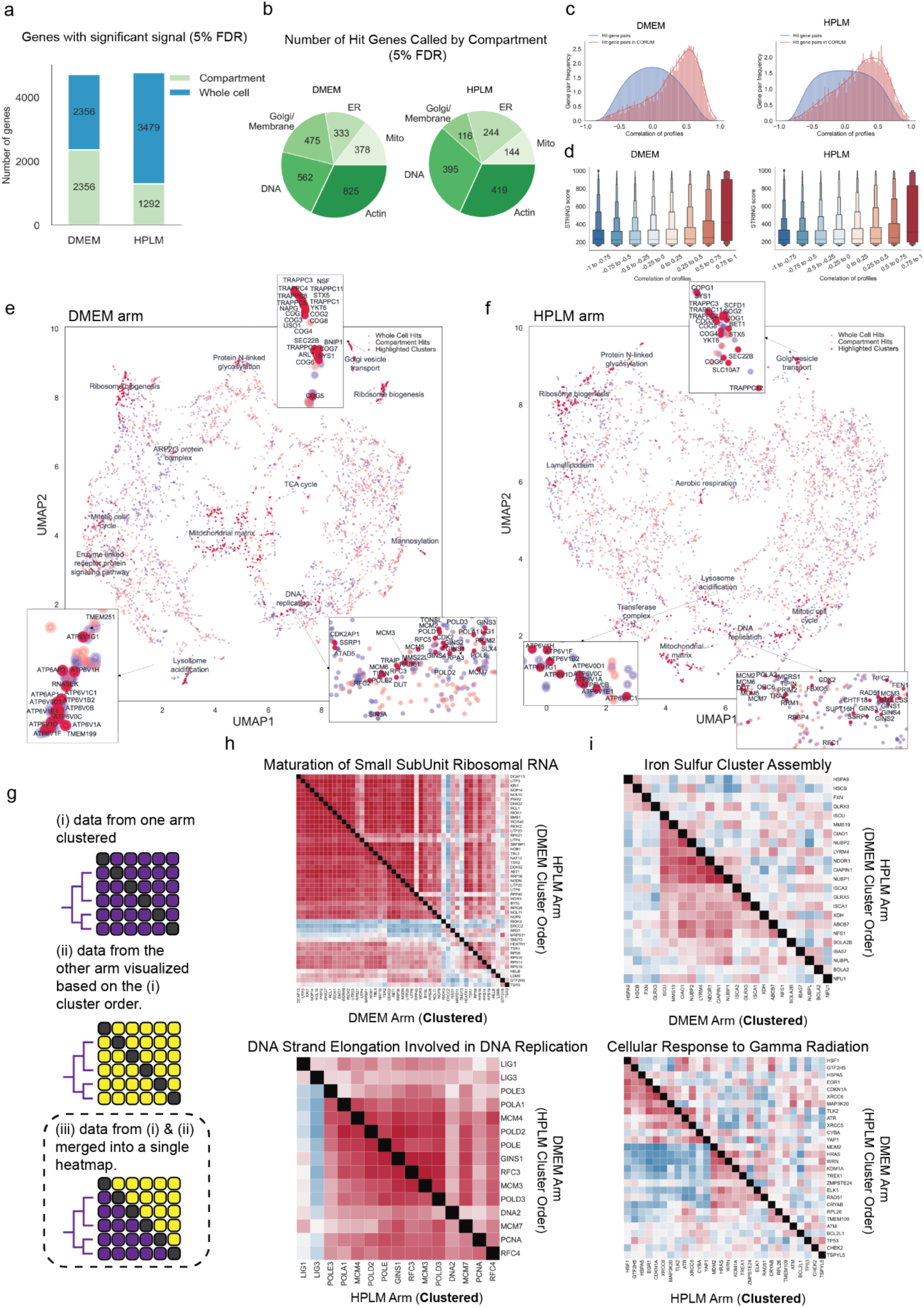
Genome-wide gene-by-environment maps. Summary of the results from two whole genome PERISCOPE screens performed in HeLa cells (DMEM and HPLM). (a) Bar graph representing the number of hit genes identified in the screen. Green represents hit genes called based on single compartments (ER, Mitochondria, Actin, DNA and Golgi/Membrane) and blue represents hit genes called based on overall gene profile. Detailed description in the methods section. (b) The distribution of hit genes called based on a single compartment (See a). It is possible for a gene to be hit in multiple compartments without being a whole cell hit, see Extended Data Fig. 3c-f for more details. (c) Distributions of optical profile correlations between random hit gene pairs versus correlations between gene pairs representing CORUM4.0 protein complexes. For the DMEM and HPLM HeLa screens, respectively, we identified 1,663 and 1,604 unique genes forming 871 and 799 clusters with at least 2/3 of components present on the profile hit list. The correlations between all possible gene pairs within a cluster (median r^DMEM^ = 0.39, r^HPLM^ = 0.27) were stronger in comparison to the background correlations between random gene pairs (median r = 0.0) (d) Boxen plot (Letter-value plot) representing STRING scores divided into bins based on PERISCOPE profile correlation between gene pairs. (e) UMAP embedding of the hit gene profiles from the DMEM HeLa dataset. Each dot represents a genetic perturbation and distance implies the correlation of profile in a two dimensional embedding. Manual annotation of cluster functions are presented for highlighted clusters based on gene ontology (GO) data sets. Example insets show coherent clustering of related genes. (f) UMAP embedding of the hit gene profiles from the DMEM HeLa dataset. Each dot represents a genetic perturbation and distance implies the correlation of profile in a two dimensional embedding. Manual annotation of cluster functions are presented for highlighted clusters based on gene ontology (GO) data sets. (g) Schematic for generation of comparative diagonally-merged heatmaps. Heatmaps display Pearson’s correlation between gene profiles from both HeLa screens and are clustered based on the data from a single screens as described in the schematic.(h, i) Heatmaps representing Pearson’s correlation between gene profiles after hierarchical clustering using Ward’s method. Gene complexes/processes were commonly enriched in both screens (h) or selectively enriched in one screen (i, the enriched screen is labeled **Clustered**) of HeLa DMEM/HPLM datasets based on the preranked GSEA analysis.

We visualized the two HeLa screens by performing 2D embedding followed by manual annotation of a subset of morphologically similar gene clusters. These clusters recapitulate known biological relationships within complexes and processes such as lysosome acidification, DNA replication, mannosylation, protein N-linked glycosylation, aerobic respiration, mitotic cell cycle, ribosome biogenesis, Golgi vesicle transport, and ARP2/3 protein complex (Fig. 3e,f). Despite differing media conditions, we found that the majority of hit genes in both screens were shared. To further visualize similarities between screens, we generated comparative diagonally-merged heatmaps (Fig. 3g) and observed that genetic perturbations identified as hits often yielded similar morphological impacts in both media types. For example, genes associated with small subunit ribosomal RNA maturation and ARP2/3 complex mediated actin nucleation, which were tightly clustered based on the DMEM profiles, exhibited strong similarity in pattern and strength of correlations in HPLM (Fig. 3h and Extended Data Fig. 8b). Likewise, perturbation profiles targeting genes involved in DNA strand elongation and PI3K-AKT-mTOR signaling, which were clustered based on the HPLM profiles, showed a similar pattern in DMEM (Fig. 3h and Extended Data Fig. 8b). These similarities in correlation patterns and strength across a variety of core processes in the same cell line indicate shared central biology and consistency of the screening method.

### Identifying media-specific perturbation signatures from optical profiles

We also identified media-specific signatures in these data, representing unique gene-by-environment interactions. To investigate these differences, we performed preranked gene set enrichment analysis (GSEA^40,41^) on both HeLa screens based on a list of morphological profiles. We quantified the strength of each profile compared to control profiles using a parameter called the “morphological signal score” (see Methods), visualizing the results in a gene enrichment map for both screens (Extended Data Fig. 8a).

Based on the GSEA analysis, 391 gene sets were enriched as hits in the DMEM screen and 321 were enriched in the HPLM screen (Supplementary Table 3). Of these, 275 were common between the two screens, 116 were specific to the DMEM screen and 46 were specific to the HPLM screen. From these, we were able to identify a subset of processes selectively enriched only in a single screen. In the DMEM screen, for example, we observed selective enrichment of processes associated with central carbon metabolism, such as NADH regeneration (a metabolic process that generates a pool of NADH by the reduction of NAD+) and glucose catabolism/glycolysis. To further investigate the enrichment of these processes in the DMEM screen, we again used comparative diagonally-merged heatmaps, observing higher levels of signal and correlation within the profiles from the DMEM screen (Fig. 3i and Extended Data Fig. 8c). Here, we also observed that Iron Sulfur cluster assembly, which is required for mitochondrial respiration^42^, and mitochondrion transcription processes were selectively enriched in the DMEM screen. Taken together, the overall enrichment of hits associated with central carbon metabolism in the DMEM screen may be reflective of metabolic differences induced by high (>25 mM) glucose levels present in DMEM^21^.

Conversely, we also observed selective enrichment of processes in the HPLM screen related to DNA damage repair, such as “cellular response to gamma radiation” or “positive regulation of DNA recombination” (Fig. 3i and Extended Data Fig. 8c). This process enrichment is also likely linked to metabolic rewiring induced by significant decreases in glucose and glutamine upon culture in HPLM, as HeLa cells have been previously shown to exhibit hallmarks of DNA damage when cultured with reduced concentrations of these nutrients^43^.

### Annotating protein complexes and cellular signaling pathways from imaging data

We next sought insights on specific structures and processes. Genes encoding various types of ribosomal proteins largely grouped into three distinct clusters (Fig. 4a). The largest cluster is enriched for genes encoding the large and the small subunits of the mitochondrial ribosome which is essential in the translation of mitochondrial genes^44^, while two other clusters show enrichment for components of the large 60S subunit and the small 40S subunit of the mature 80S eukaryotic ribosome^45^. This example highlights the ability of optical pooled screens to capture structural information, as recently demonstrated^12^. We also found that signaling pathways were often well-captured: as an example, perturbations targeting the phosphatidylinositol 3-kinase/AKT serine-threonine protein kinase (PI3K/AKT) signaling pathway largely fall into two distinct clusters (Fig. 4b). This pathway is involved in the cell cycle, growth and proliferation, and implicated in the progression of various cancers.^46^ Interestingly, components which have a stimulatory effect on the pathway like RPTOR, MTOR or MYC strongly correlate with each other and also demonstrate a significant anti-correlation to inhibitory factors such as PTEN, TSC1 or TSC2. The ability of morphological profiles to distinguish the directionality of these signaling factors is a useful tool in understanding the underlying biology.

**Figure 4.**
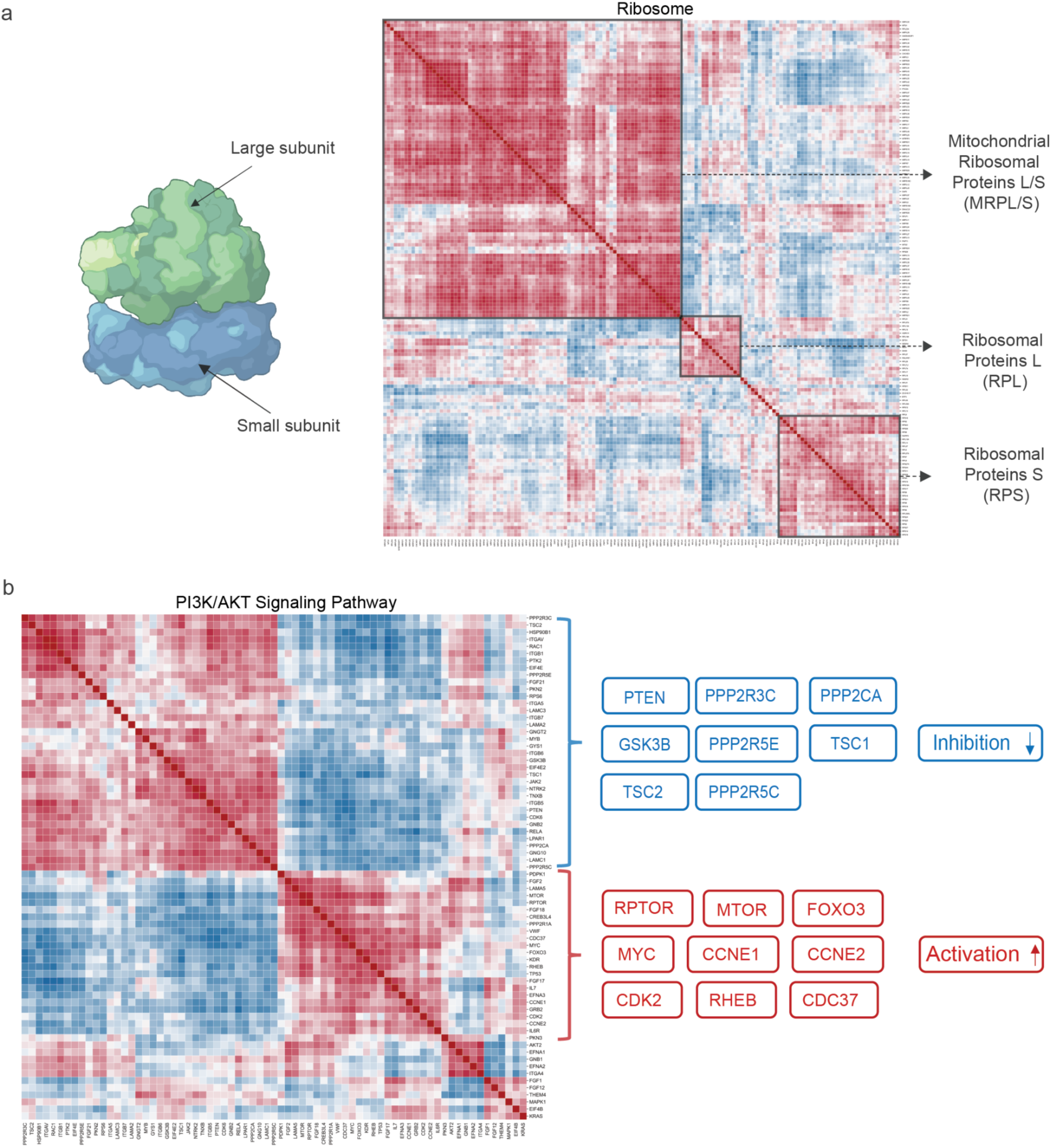
Clustering by optical profiles captures physical interactions and signaling pathway relationships. (a) Heatmaps representing Pearson’s correlation between gene profiles after hierarchical clustering using Ward’s method. The ribosomal gene complex was enriched in the DMEM HeLa dataset based on the preranked GSEA analysis. Different subset of ribosomal genes enriched in certain clusters are highlighted in the heatmap. Ribosome image created with Biorender. (b) Heatmaps representing Pearson’s correlation between gene profiles after hierarchical clustering using Ward’s method. The genes associated with the PI3K/AKT signaling pathway were enriched in the DMEM HeLa dataset based on the preranked GSEA analysis. The table on the right highlights the genes with activatory/inhibitory effects in accordance with the correlation/anti-correlation between profiles from the heatmap.

### Genome-wide screens for subcellular phenotypes of interest

High-dimensional profiles generated by PERISCOPE are composed of thousands of individual phenotypic features. Having seen that full morphological profiles capture biologically meaningful patterns of similarity, we next explored whether the datasets could be used to conduct genome-wide screens for individual morphological phenotypes of interest. To explore the single-feature screen space, we looked at each feature in our feature-selected datasets and generated a most-perturbed gene list and then assayed for GO enrichment within that list. Features with GO enrichment were fairly evenly distributed across imaging channels for both HeLa screens (Fig. 5a) which is unsurprising given that each channel contributed similarly to our profile-based hit lists (Figures 2b and 3b) and all canonical channels contribute fairly evenly to profile strength in the Cell Painting Assay^47^. Features with GO enrichment were not as evenly distributed across feature classes (Figure 5b); the feature class with the largest number of features - texture - also had the largest fraction of its features (37%: 370 out of 1001 features) enriched for a GO term, although it is likely that many such features correlate to some degree - our feature selection step removes only the most highly correlated features. Notably, AreaShape and Intensity features, which are often emphasized in other studies because of their biological interpretability, were less specifically enriched than less readily-understandable categories such as Texture and Correlation.

**Figure 5.**
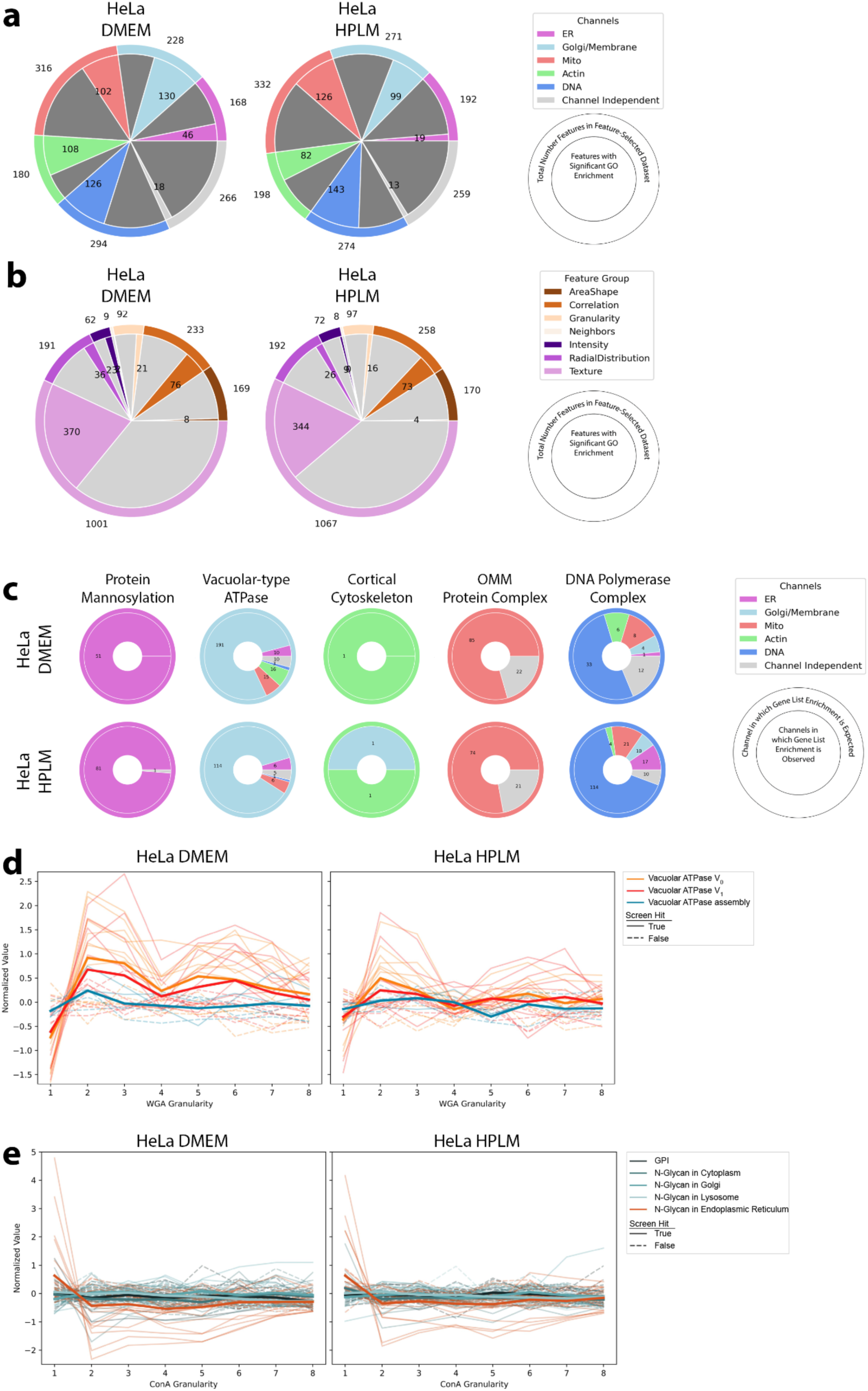
Identifying biological pathways using individual subcellular image features in HeLa datasets. (a) GO enrichment is found in many individual features in a manner that is fairly evenly distributed across the cellular structures (i.e. channels) imaged in PERISCOPE. Outer ring is the total number of features in our feature-selected dataset. Inner ring is the number of features that show GO enrichment. (b) GO enrichment in individual features is not distributed evenly across classes of features. Outer ring is the total number of features in our feature-selected dataset. Inner ring is the number of features that show GO enrichment. (c) Given gene groups whose protein products are expected to function specifically in a cellular structure imaged in PERISCOPE, are specifically enriched in hit lists for features in those compartments. Outer ring indicates the channel in which enrichment is expected. Inner ring is the breakdown of actual channels that show enrichment for the gene group. (d) Disruption of the Vacuolar ATPase (either V^0^ or V^1^ subunit) but not genes involved in its assembly causes a decrease in WGA signal in small structures as seen specifically with screen feature WGA_Granularity_1 but not larger granularities. Each trace is a single gene; those genes that are not hits in the screen are dashed. Bold lines are the mean of all genes in the group. (e) Loss of function in genes involved in N-Glycan synthesis in the Endoplasmic Reticulum but not in other organelles nor the GPI synthesis pathway causes an increase of ConA signal in small structures as seen specifically with screen feature ConA_Granularity_1 but not larger granularities.

To support the validity of the dataset’s single feature screens, we looked at groups of genes whose protein products are known to function in the compartments that we labeled in PERISCOPE and determined which features had hit lists that were enriched for those groups. In Figure 2c we showed that perturbing these groups of genes produces signal across the channels whereas we show in Figure 5c that there is specific enrichment in our hit lists for features in expected categories for protein mannosylation, vacuolar-type ATPase, cortical cytoskeleton, and outer mitochondrial membrane protein complex, though unsurprisingly perturbation of DNA polymerase generated a more pleiotropic phenotype.

As a complementary way to assess specificity, we focused on perturbations that altered granularity features^48^, a measure of the signal present within differently-sized intracellular structures, from small cellular details to increasingly large scale structures, relative to the total signal. We found that disruption of the vacuolar ATPase (either V_0_ or V_1_ subunit), but not genes involved in its assembly, causes a decrease in WGA signal in small structures (Figure 5d, visualized in Extended Data Figure 9f) for both HeLa datasets. We also found that knocking out genes involved in N-Glycan synthesis in the Endoplasmic Reticulum but not in other organelles nor the GPI synthesis pathway causes an increase of ConA signal in small structures (Figure 5e, visualized in Extended Data Figure 9g) in our HeLa datasets. The A549 screen did not show as much specific enrichment (Extended Data Figure 9a-c) but nevertheless still also showed a specific vacuolar ATPase phenotype (Extended Data Figure 9d).

Each screen dataset includes 3,973 features from our adapted Cell Painting assay. Although there is significant redundancy, particularly among texture and granularity metrics, there are nevertheless dozens of morphological phenotypes of interest to the biological community whose genome-wide screening data can now be explored and hits pursued.

A primary advantage of image-based profiling over traditional microscopy is that the former allows quantitative and automated assessment of phenotypic features, overcoming the subjectivity of analyzing images by eye. Nonetheless, our atlas contains over 30 million individual cell images that can be evaluated for phenotypes of interest by a trained eye. Thus, to enhance the usefulness of these datasets, we have developed an atlas cell retrieval tool (see Methods) enabling the retrieval of individual images of cells containing perturbations of interest (Extended Data Fig. 5a-f). Using this tool, we show that it is possible to find examples of readily interpretable image-based phenotypes, such as the depletion of TOMM20 signal in cells containing sgRNAs targeting TOMM20 (Extended Data Fig. 5e), but also that most single gene knockout phenotypes are more subtle (Extended Data Fig. 5b-d,f), demonstrating the usefulness of computational feature extraction and profiling beyond simple visual inspection.

### Identification of TMEM251/LYSET as a Golgi-resident protein essential for mannose-6-phosphate-dependent trafficking of lysosomal enzymes

Having observed that genes cluster by function using morphological profiles, we next sought to ascertain the function of uncharacterized genes based on profile similarity. We focused on the poorly characterized gene TMEM251, which clustered with genes involved in lysosomal acidification (Fig. 3e). GSEA of genes ranked by similarity to the TMEM251 KO profile in the HeLa DMEM dataset revealed enrichment for V-ATPase subunits and Golgi components, especially those related to glycosylation (Fig. 6a,b). Based on these term enrichments, we compared the subcellular localization of TMEM251 relative to the Golgi and lysosomes in HT1080 cells, which were selected for their relative TMEM251 growth dependency^31^. TMEM251 localized primarily to the Golgi, with negligible localization to lysosomes (Fig. 6c). TMEM251 knockdown (KD) with CRISPRi created strong phenotypes in the WGA channel (Fig 6d), contributed by a striking accumulation of WGA fluorescence in LAMP1-positive lysosomes (Fig. 6d). This phenotype was seen for most of the perturbations bearing strong profile similarity to TMEM251 with the notable exception of SLC35A2, which was the most similar gene to TMEM251 at the profile-level (Fig. 6e and Extended Data Fig. 10a).

**Figure 6.**
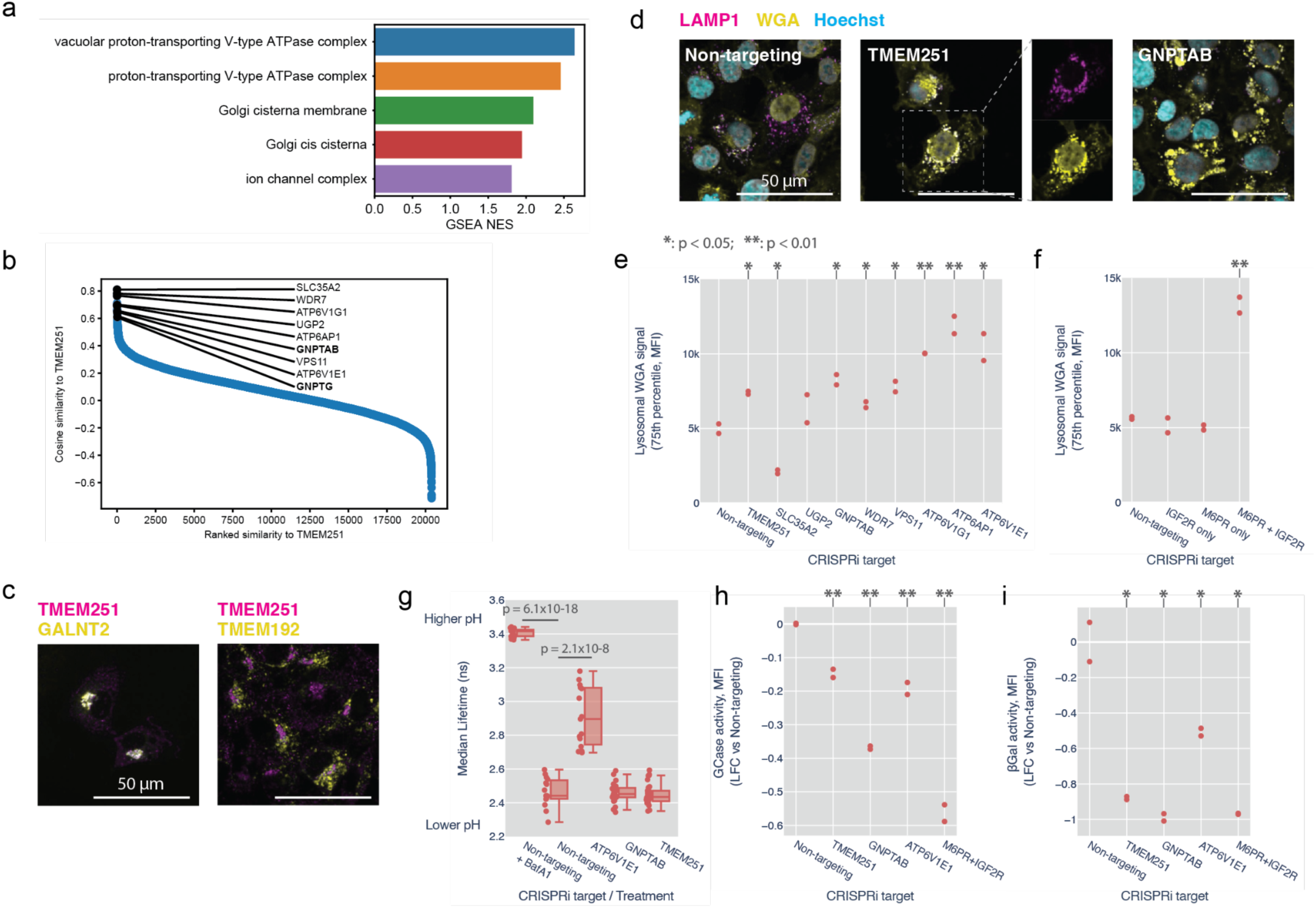
TMEM251 is essential for M6P-dependent trafficking of lysosomal enzymes. (a) GSEA of genes preranked by cosine similarity to TMEM251 KO morphology. (b) Waterfall plot of the distribution of cosine similarities to TMEM251 morphology. Representative genes involved in glycosylation, trafficking and lysosomal acidification are highlighted. (c) TMEM251 localization was examined by confocal imaging of cells expressing fluorescent reporter of either GALNT2 (Golgi) or TMEM192 (lysosome) and stained for TMEM251. (d) Confocal images of cells with knockdown of genes indicated, co-stained with WGA and LAMP1 antibody. See supplementary figure for other perturbations. (e-f) Quantification of lysosomal WGA staining after CRISPRi knockdown of the indicated genes. Plotted are the upper quartiles of median per-cell lysosomal WGA intensity in biological replicates. (g) Boxplot of LAMP1-mScarlet fluorescence lifetimes, which correlates with lysosomal pH, for the indicated perturbations. Each point represents the median lifetime of lysosomal fluorescence in an image (n≥15 per condition). (h-i) log10 fold-changes of glucosylceramidase and beta galactosidase activity relative to non-targeting controls for the indicated CRISPRi knockdowns. Each point represents the median of per-cell total fluorescence intensity (MFI) in biological replicates, relative to non-targeting controls. (Statistical analysis: 2-tailed t-test vs Non-targeting for e to i)

How could a Golgi-resident protein influence glycan storage in the lysosome? We postulated that the lysosomal WGA phenotype was due to impaired biogenesis of lysosomal proteins in the Golgi. Notably, GNPTAB/GPNTG showed strong phenotypic similarity to TMEM251 in PERISCOPE and human LOF of TMEM251 results in a clinical presentation similar to that of human LOF in GNPTAB/GNPTG^49^. We therefore hypothesized that TMEM251 may participate in the mannose-6-phosphate (M6P) pathway. In this pathway, N-Acetylglucosamine-1-Phosphate Transferase (encoded by GNPTAB) attaches a phospho-GlcNac from UDP-GlcNac onto a terminal mannose that ultimately forms M6P^50^. M6P is recognized by either of two receptors, M6PR and IGF2R, and released in the lysosome in a pH-dependent manner. To further corroborate this hypothesis, we compared the phenotype of cells singly or doubly perturbed for M6PR and IGF2R. In double knockdown cells we observed a significant increase in lysosomal WGA accumulation, whereas single knockdowns were indistinguishable from wildtype, consistent with the primary screen (Fig. 6f and Extended Data Fig. 10b).

Because of the strong morphological similarity between TMEM251 and V-ATPase subunits, we examined the effect of TMEM251 KD on lysosomal pH using a fluorescence lifetime sensor^51^. Whereas treatment with Bafilomycin A1 or ATP6V1E1 KD robustly alkalinized lysosomes, neither GNPTAB nor TMEM251 KD significantly changed lysosomal pH (Fig. 6g and Extended Data Fig. 10c). We therefore reasoned that acidic lysosomal pH might be required for proper trafficking and functioning of lysosomal enzymes downstream of TMEM251’s Golgi function and that the optical profile induced by V-ATPase perturbation is dominated by this function. We tested the activity of two lysosomal enzymes that require M6PR for proper localization. Glucosyl cerebrosidase is recognized by SCARB2, which in turn interacts with M6PR to traffic to the lysosome^52^. TMEM251, GNPTAB, ATP6V1E1 and the M6PR/IGF2R double KDs all reduced Glucosyl cerebrosidase activity (Fig. 6h and Extended Data Fig. 10d). Beta galactosidase activity was even more dramatically impaired by these KDs (Fig. 6i and Extended Data Fig. 10e). During preparation of this manuscript, two independent groups reported the function of TMEM251 in the biogenesis of M6P and renamed the protein LYSET ^53,54^. Our results independently support and validate a role for TMEM251 in lysosomal protein trafficking through the M6P-system.

## Discussion

Pooled optical screens are a powerful new approach for generating high-dimensional genotype-phenotype maps with single cell resolution. Our studies demonstrate that these maps can now be generated routinely at scale, enabling the interrogation of genome-scale perturbation effects using standard laboratory equipment (a widefield fluorescence microscope) and scalable, distributed open source analysis pipelines. Further, the cost is very low per cell profile: ∼$0.001/cell for the described datasets. We believe this combination of accessibility and cost effectiveness make PERISCOPE-style screens a democratizing platform technology for linking genotypes to cellular programs.

In addition to being practical, PERISCOPE generates rich, data-driven representations of gene function. A central goal of massively parallel genetic screens is to understand how genes coordinate to produce complex cell phenotypes and in this regard PERISCOPE is valuable both as a profiling technology – generating high-dimensional representations of cell state – and as highly-parallelized screens of subcellular biological parameters (e.g. cell size, organelle size/shape/number). We demonstrate that whole-cell optical profiles can be used to reconstruct relationships between genes in biological pathways and proteins in complexes. We further demonstrate that spatially restricted subcellular phenotypes in these data can be used to gain mechanistic insight into gene function, as in the case of Golgi-specific phenotypes affected by perturbation of TMEM251. Further, we found that individual morphological features (e.g. regional granularity) could also be used to classify genes by function such as with genes involved in V-ATPase assembly or N-glycan synthesis.

Massively parallel CRISPR modifier screens have been proven to be very useful for mapping gene-by-environment interactions at scale. By enabling facile, cost effective genome-scale screening with high-dimensional cell profiling, we demonstrate that genetic perturbations can be readily combined with environmental perturbations to produce rich, high-resolution maps to systematically interrogate gene-by-environment interactions at genome scale. As an example, we show how such maps can uncover media-specific effects on cellular programs, but we additionally envision using this platform to execute genome-wide screens for modifiers of therapeutic compound-induced phenotypes, or to carry out genetically anchored CRISPR screens^55^ to elucidate genetic interaction networks.

Beyond the current scope, there are several improvements that could be built upon the foundation of the work presented here. In its current form, the PERISCOPE platform could be deployed to explore the effects of other CRISPR-based perturbations such as CRISPR-a^56,57^, CRISPR-i^58,59^, or base editing^60–62^, where sgRNAs can be expressed as a DNA Pol II transcript (as in CROP-seq). In this study, we profile two cancer cell lines, HeLa and A549, but our pipelines are amenable to screening a wide variety of 2D cell models, including cell lines and primary cells, though assay scale and data quality are cell density-dependent. Our screens demonstrate that significant signal is present in every measured cell compartment, and highly multiplexed imaging technologies such as CODEX^63^ and CyCIF^64^ could improve the sensitivity and robustness of PERISCOPE by capturing a wider range of perturbation effects or enabling the inclusion of ground truth epitopes to anchor biological interpretation. Extracting biological signals from fluorescence multicolor images is a compelling machine learning problem which will likely be improved using various forms of deep learning, such as self-supervised learning, to extract features. Though such features lack inherent interpretability, they are more powerful for capturing similarities and can be useful for many applications^65,66^.

In sum, this study lays the groundwork for building high-dimensional morphology-based perturbation maps at scale and presents the first genome scale atlas of human cell morphology. Containing more than 30 million perturbation-assigned cell images, this atlas is a useful resource for biological interrogation as well as for the development and testing of new computational image analysis methods. All data and analysis tools are open source and freely available (see Code and Data availability).

## Methods

### Library design

The whole genome library was designed to target 20,393 genes with ∼4 sgRNAs per gene for a total of 80,408 sgRNAs. 47,792 sgRNAs were selected from the Brunello CRISPR library (Addgene #73179), 20,520 sgRNAs were selected from the TKO V3 CRISPR library (Addgene #90294), and 12,096 sgRNAs were selected from the extended CRISPR library published by the Broad Institute’s Genetic Perturbation Platform (Addgene #73178). Additionally, 601 non-targeting sgRNAs were included as negative controls. All sgRNA sequences were selected/designed to maintain a balanced nucleotide distribution at each base position, which facilitates optical barcode calling. The CRISPR library was designed for complete library deconvolution with eleven bases and for levenshtein error correction with twelve bases.

### Library cloning

In order to prepare pooled plasmid libraries, targeting and non-targeting guide subpools were first individually amplified by dialout PCR using orthogonal primer pairs.^67^. PCR products were purified using the QIAquick PCR Purification Kit (Qiagen LLC #28104). The amplified libraries were cloned into the CROPseq vector (Addgene #86708) via Golden Gate assembly using BsmBI restriction sites as previously described^15^. To prevent self ligation events in Golden Gate reactions, the CROPseq vector was pre-digested and purified via gel extraction using the QIAquick Gel Extraction Kit (Qiagen LLC #28706) in order to remove the filler sequence. The resulting plasmid libraries were purified and concentrated via SPRI bead cleanup before being transformed into electrocompetent cells (Lucigen Endura, VWR International LLC #71003-038) for plasmid library amplification. Following transformation, bacterial cells were grown in liquid cultures for 18 hours at 30°C before extracting the plasmid DNA. The plasmid library was validated via Next Generation Sequencing as described in NGS methods below.

### Tissue culture

A549 cells were cultured in High-glucose DMEM (VWR International LL #45000-304) supplemented with 2mM L-glutamine (Life Technologies Corporation #25030081), 100 U/mL penicillin-streptomycin (Life Technologies Corporation #15140163), and 10% heat-inactivated fetal bovine serum (Sigma-Aldrich Inc #F4135-500ML). HEK293FT cells were cultured in DMEM-GlutaMax, pyruvate (Thermo Fisher Scientific #10569010) supplemented with 10% heat-inactivated fetal bovine serum, and 100 U/mL penicillin-streptomycin, and 2mM L-glutamine. HEK293FT cells were also cultured without antibiotics 24 hours prior to lentiviral packaging. HeLa cells in the conventional media screen were cultured in DMEM (VWR International LL #45000-304) supplemented with 10% dialyzed FBS (ThermoFisher Scientific #26400044). HeLa cells in the physiological media screen were cultured in HPLM (Thermo Fisher Scientific #A4899101) supplemented with 10% dialyzed FBS.

### Lentivirus production

Prior to lentivirus production, the plasmid pools for targeting and nontargeting sgRNAs were combined resulting in a 10% (m/m) of nontargeting sgRNAs and a 90% (m/m) of targeting sgRNAs. 24 hours before transfection, HEK293FT cells were seeded on 10cm2 dishes at a density of 100,000 cells/cm2 using antibiotic free media. Lentivirus was generated using the Lipofectamine 3000 (Thermo Fisher Scientific L3000015) transfection kit and packaging plasmids pMD2.G (Addgene #12259) and psPAX2 (Addgene #12260). HEK293FT cells were transfected with a plasmid ratio of 2:3:4 (by mass) of pMD2G, psPAX2, and plasmid library, respectively. Media was exchanged four hours after transfection. Lentivirus was harvested 48 hours after media exchange and filtered through a 0.45um cellulose acetate filter (Corning 431220). The viral supernatant was incubated in dry ice until frozen and stored at -80°C.

### Lentivirus titering

A viral titer was individually determined for A549 and HeLa cells. A549 cells were seeded at a density of 100,000 cells/cm^2^ while HeLa cells were seeded at a density of 150,000 cells/cm in a 6 well format. The seeded cells were transduced with the viral library by supplementing their media with 8 μg/mL of polybrene (Sigma-Aldrich Inc # TR-1003) and adding a variety of viral volumes ranging from 0 μL to 50 μL prior to centrifugation at 1000 g for 2 hours at 33°C. After centrifugation, the cells were incubated at 37 °C for 4 hours followed by a media exchange. At 24 hours post-infection, cells were divided into media containing either 0 μg/mL or 2 μg/mL of puromycin (Life Technologies #A1113803). Cells in both media conditions were incubated at 37°C for 72 hours. Following incubation, cells were counted and multiplicity of infection (MOI) was estimated by the ratio of surviving cells in the 2 μg/mL puromycin conditions over puromycin free conditions. Infectious units per microliter (ifu/μL) were then calculated by multiplying the MOI by the original cell seeding density and dividing by the viral volume added. The ifu/μL for each viral volume were averaged and used to estimate viral volume required to achieve an MOI between 0.1 and 0.3.

### Lentivirus transduction

For screens, cells were transduced with the genome-wide viral library in a 6-well format by adding 8 μg/mL of polybrene and the volume of viral supernatant calculated for an MOI of 0.2 as well as a non-infection control with 0 μL of viral supernatant. Cells were centrifuged at 1000 g for 2 hours at 33°C. At 4 hours post-infection, media was exchanged. At 24 hours post-infection, the infected cells were passaged into T-225 flasks (VWR International LLC #47743-882) containing media supplemented with 2 μg/mL puromycin. A fixed number of cells (∼300,000) for the infection and uninfected conditions were set aside and seeded in a 6-well plate format under media containing either 0 μg/mL or 2 μg/mL of puromycin. All cells were incubated at 37°C for 72 hours. Following the 72 hours of selection, the cells seeded in the 6-well plate were counted and the MOI was calculated as described above.

### A549 screen

A549-TetR-Cas9 cells were transduced with the genome-wide viral library in three biological replicates by seeding cells at a density of 150,000 cells/cm2 in a 6-well format and performing lentiviral transduction as described above. A total of 240,000,000 cells were transduced at an MOI of 0.2 for a cell library representation of 300 cells/sgRNA post transduction. After antibiotic selection, the cells were cultured in conventional DMEM media for two days. Prior to induction of Cas9 expression, a sample of 25,000,000 cells per biological replicate were lysed and prepared for Next Generation Sequencing (NGS) as described below. These samples were used to confirm the target representation. Cas9 expression was induced with 2 μg/mL doxycycline spiked in conventional DMEM media. Throughout Cas9 expression, cells were cultured in T-225 flasks and passaged once the flasks reached 70% confluency. Between passages, a minimum of 24,000,000 cells were re-seeded per biological replicate thus maintaining a representation of 300 cells/sgRNA. The cells were supplemented with 2 μg/mL of doxycycline every two days by exchanging the culturing media. On day 5 of Cas9 expression, the cells were seeded into nine 6-well glass-bottom plates (Cellvis #P06-1.5H-N) at a density of 19,800 cells/cm. A total of 13,000,000 cells across the three biological replicates were seeded in optical plates with the expectation that cell populations will double at least once before fixation. The remainder of the cells were kept in T225 flasks and cultured until day 7 of Cas9 expression where a sample of 13,500,000 cells from each biological replicate were lysed and prepared for NGS analysis. This analysis was then used to determine sgRNA dropout rates due to lethal CRISPR events. 48 hours after being seeded in optical plates, the cells were fixed with 4% paraformaldehyde in 1X PBS for 30 minutes, followed by *in situ* sequencing (ISS) as described below. After RCA amplification in ISS, the cells were stained with cell compartment-specific probes as described in Cell Staining and phenotypic images were acquired. The disulfide-linked probes were de-stained by cleaving the disulfide bridge between the probe and its fluorophore with 50 mM TCEP (Thermo Fisher Scientific #363830100) in 2X saline-sodium citrate (SSC) for 45 minutes at room temperature. After destaining phenotypic probes, the cells are washed three times with 1X PBS-T (1X PBS + 0.05% Tween-20) before performing 12 cycles of *in situ* sequencing by synthesis (ISS).

### HeLa screens

HeLa-TetR-Cas9 were transduced with the genome-wide viral library in three biological replicates by seeding cells at a density of 210,000 cells/cm^2^ in a 6-well format and performing lentiviral transduction as described above. A total of 240,000,000 cells were transduced at an MOI of 0.2 for a cell library representation of 300 cells/sgRNA post transduction. After antibiotic selection, the transduced cells were cultured in conventional DMEM media until a representation of 600 cells/sgRNA was achieved. In order to confirm the target representation, a sample of 20,000,000 cells from each biological replicate were lysed and prepared for Next Generation Sequencing (NGS) as described below. The cell library was then divided into two culturing conditions, conventional DMEM media and Physiological HPLM media (media formulations are described above). Simultaneous to the addition of these two media conditions, Cas9 expression was induced with 2 μg/mL doxycycline (reagent reference) for 7 days. Throughout Cas9 expression, cells for each condition were cultured in T-225 flasks and passaged once the flasks reached 70% confluency. Between passages, a minimum of 24,000,000 cells were re-seeded per biological replicate thus maintaining a representation of 300 cells/sgRNA for each media condition. The cells were supplemented with 2 μg/mL of doxycycline every two days by exchanging the culturing media. On day 5 of Cas9 expression, the cell libraries under both media conditions were seeded into five 6-well glass-bottom plates (Cellvis #P06-1.5H-N) at a density of 42,000 cells/cm2. A total of 14,000,000 cells across the three biological replicates were seeded in optical plates for each media condition with the expectation that cell populations will double at least once before fixation. The remainder of the cells were kept in T225 flasks and cultured until day 7 of Cas9 expression where a sample of 20,000,000 cells from each biological replicate were lysed and prepared for NGS analysis. This analysis was then used to determine sgRNA dropout rates due to lethal CRISPR events. 48 hours after being seeded in optical plates, the cells were fixed with 4% paraformaldehyde in 1X PBS for 30 minutes, followed by *in situ* sequencing (ISS) as described below. After RCA amplification in ISS, the cells were stained with cell compartment-specific probes as described in Cell Staining and phenotypic images were acquired. The disulfide-linked phenotypic probes were destained by cleaving the disulfide bridge between the probe and its fluorophore with 50 mM TCEP (Thermo Fisher Scientific #363830100) in 2X saline-sodium citrate (SSC) for 45 minutes at room temperature. After probe destaining, the cells are washed three times with 1X PBS-T (1X PBS + 0.05% Tween-20) before performing 12 cycles of ISS.

### Synthesis of de-stainable phenotyping probes

Due to the spectral overlap between the fluorescent dNTPs required for ISS and the available fluorophores for phenotypic markers, the probes used to label the mitochondria and the endoplasmic reticulum (ER) were synthesized in house to include a disulfide bridge between the probe and its fluorophore that will allow for cleavage of the fluorophore after imaging. For mitochondria labeling, the secondary anti-TOMM20 antibody, F(ab’)2-goat-anti-rabbit IgG (H+L) (Thermo Fisher #31239) was conjugated to Alexa Fluor 594-Azide (Thermo Fisher #A10270). For ER labeling, the protein Concanavalin A (ConA) (Sigma Aldrich #C2010) was conjugated to Cyanine 5-Azide (Lumiprobe #B3030). In the synthesis of these probes, we leveraged the thermal stability and high specificity of the click chemistry reaction between dibenzocyclooctyne (DBCO) and Azide groups. Hence, the anti-TOMM20 antibody and the ConA protein were functionalized for click chemistry with the addition of a NHS-SS-DBCO (DBCO) (Sigma Aldrich #761532) molecule that subsequently reacted with the azide groups linked to their respective fluorophores. Prior to functionalizing the probes, the anti-TOMM20 antibody and ConA protein were diluted to 1.1 mg/mL and 2 mg/mL in freshly prepared 0.1M sodium phosphate solutions at pH 8.5 and 6.8 respectively. The DBCO was freshly dissolved to 10mg/mL in anhydrous DMSO (Sigma Aldrich #227056). The diluted proteins and DBCO were then incubated for 2 hours at 4°C while shaking. The molar ratios and buffers for this reaction are listed in Table. 1. Following incubation, the reaction was quenched with 2M Tris HCl (pH 7.4) at a 10% reaction volume. The resulting product was purified using Zeba columns (Thermo Fisher #89883). Product retention after column purification was ∼90%. The azide linked fluorophores were diluted to 10mg/mL in anhydrous DMSO and reacted with their respective functionalized probes at 3:1 molar ratio. This reaction proceeded for 20 hours at 4°C while shaking; reaction vials were protected from light during this incubation. The final product was purified by running each reaction through three Zeba columns in order to do a final buffer exchange into 1X PBS. After synthesis the destainable probes were stored at -20°C.

**Table 1.**
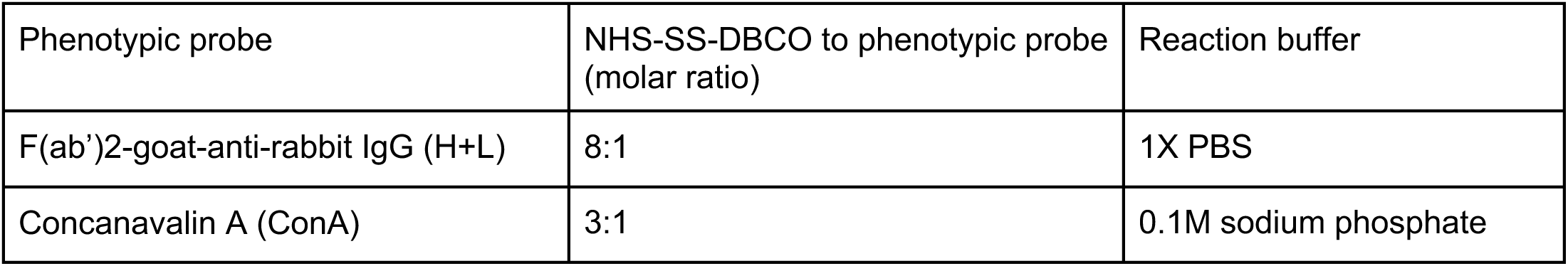
Functionalization of phenotypic probes for click chemistry based synthesis of destainable cell markers.

| Phenotypic probe | NHS-SS-DBCO to phenotypic probe<br>(molar ratio) | Reaction buffer |
| --- | --- | --- |
| F(ab')<sub>2</sub>-goat-anti-rabbit IgG (H+L) | 8:1 | 1X PBS |
| Concanavalin A (ConA) | 3:1 | 0.1M sodium phosphate |

### *In situ* sequencing

The in situ sequencing of sgRNAs required three enzymatic steps, a targeted reverse transcription of the sgRNA (RT), the formation of a circular DNA template (Gap-fill and ligation), and the amplification of that template through rolling circle amplification (RCA). Prior to the enzymatic reactions, cells were fixed with 4% paraformaldehyde (Electron Microscopy Sciences #15714) in 1X PBS for 30 minutes at room temperature and then permeabilized with 70% ethanol (VWR International #76212-358) for 30 minutes at room temperature. To prevent sample dehydration after permeabilization, the ethanol was removed over six serial dilutions with PBS-T (1X PBS + 0.05% Tween-20). After permeabilization, the RT solution was prepared according to the table below and applied to the cells. Cells in the RT solution were incubated at 37 °C overnight.

**Table 2.**
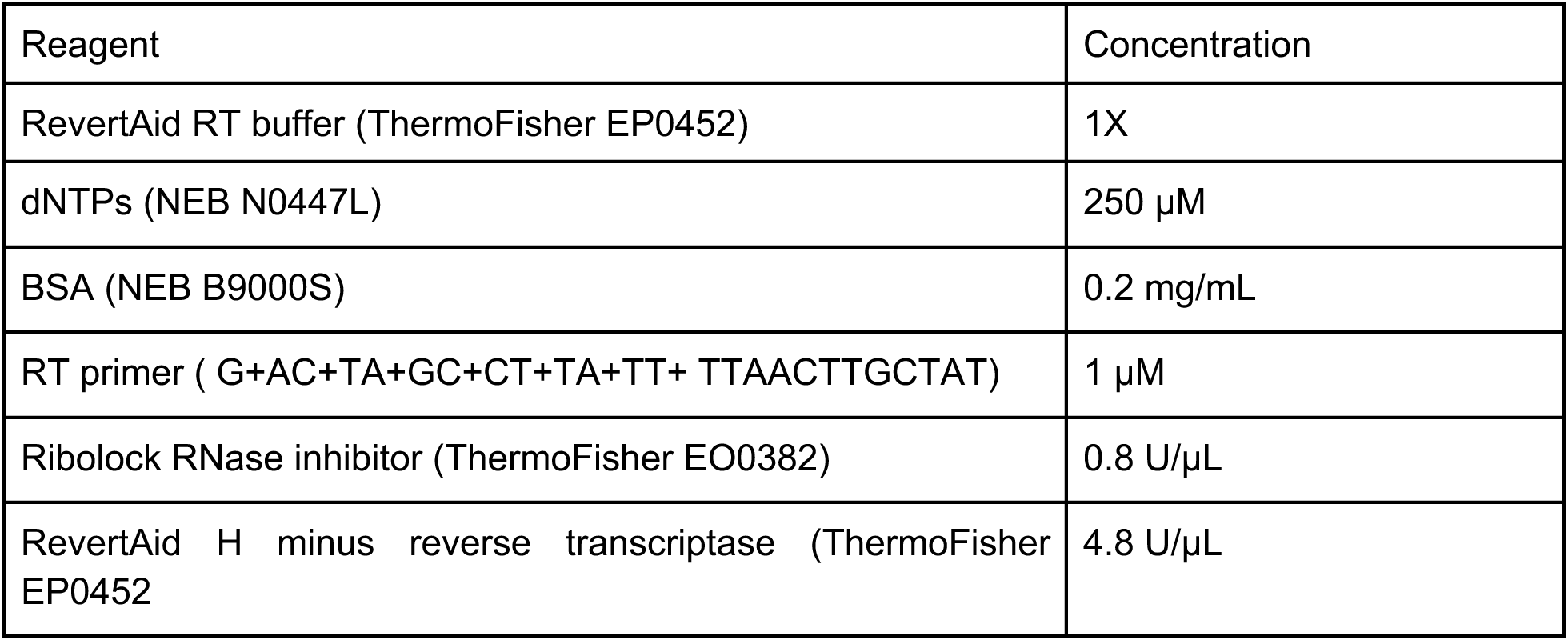
Reverse Transcription (RT) Solution.

Following reverse transcription, the cells were washed five times with PBS-T and post-fixed with 3% paraformaldehyde and 0.1% glutaraldehyde (Electron Microscopy Sciences #16120) in 1X PBS for 30 minutes at room temperature. After post-fixation, the cells were washed three times with PBS-T. The gap-fill and ligation solution was then prepared according to the table below and then added to the cells, which were then incubated at 37 °C for five minutes and 45 °C for 90 minutes.

**Table 3.**
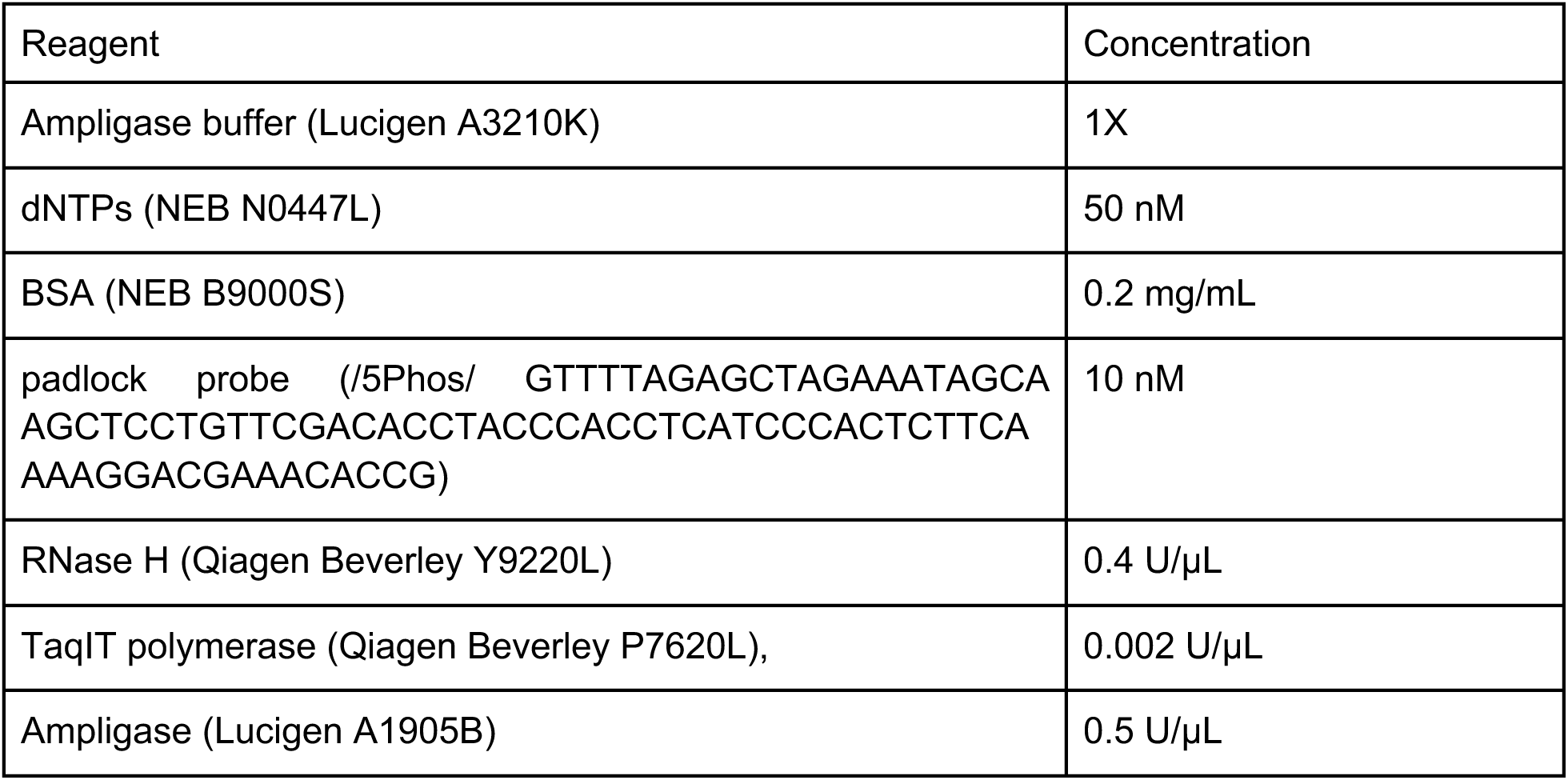
Gap-fill and Ligation Solution.

After gap-fill and ligation, the cells were washed three times with PBS-T. The RCA solution was then prepared according to the table below. The cells in the RCA solution were incubated at 30°C overnight. Following incubation, the cells were washed three times with PBS-T.

**Table 4.** Rolling Circle Amplification (RCA) Solution.

| Reagent | Concentration |
| --- | --- |
| Phi29 buffer (ThermoFisher EP0094) | 1X |
| dNTPs (NEB N0447L) | 250 µM |
| BSA (NEB B9000S) | 0.2 mg/mL |
| Glycerol | 5% |
| Phi29 DNA polymerase (ThermoFisher EP0094) | 1 U/µL |

### Phenotypic labeling

After rolling circle amplification, the cells were prepared for phenotypic labeling by incubating them with a blocking buffer containing 1% BSA (Seracare Life Sciences #1900-0016) in 1X PBS for 10 minutes at room temperature. After blocking, a primary staining solution containing rabbit Anti-TOMM20 antibody (Abcam #ab78547), Alexa Fluor 488 Phalloidin (ThermoFisher #A12379), ConA-SS-A647, and WGA-A750 (WGA protein by Vector Labs #L-1020-20, custom conjugation to A750 fluorophore by Arvys Proteins) was prepared in 1X PBS and applied to the cells for 45 minutes at room temperature. Following incubation with the primary staining solution, the cells were washed three times with 1X PBS-T and a secondary staining solution containing F(ab’)2-goat-anti-rabbit IgG (H+L)-SS-A594 was prepared in blocking buffer and applied to the cells for 30 minutes at room temperature. The phenotypic probes for the primary and secondary staining solutions were diluted according to the dilution factors listed in Table 5. Dilution factors for each probe were determined prior to screening by doing a serial titration of individual stains. After incubation with the secondary staining solution, the cells were washed with 1X PBS-T three times allowing the plate to sit at room temperature for 5 minutes between washes. Finally, the cells were placed in a freshly prepared DAPI staining solution containing 200 ng/mL DAPI (Sigma-Aldrich #D9542-10MG) diluted in 2X SSC. The cells were incubated in the DAPI staining solution for 10 minutes at room temperature prior to imaging.

**Table 5.** Dilution factors of phenotypic probes.

| Phenotypic probe | Staining solution | A549-TetR-Cas9<br>Dilution factor<br>or<br>concentration | HeLa-TetR-Cas9 cells<br>Dilution factor or concentration |
| --- | --- | --- | --- |
| Alexa Fluor 488 Phalloidin | Primary Staining Solution | 1:500 | 1:200 |
| Rabbit Anti-TOMM20 antibody | Primary Staining Solution | 1:1000 | 1:2000 |
| WGA-A750 | Primary Staining Solution | 1:1000 | 1:1600 |
| ConA-SS-A647 | Primary Staining Solution | 1:400 | 1:800 |
| F(ab')<sub>2</sub>-goat-anti-rabbit IgG (H+L)-SS-A594 | Secondary Staining Solution | 1:600 | 1:400 |

### Sequencing by synthesis

After destaining the phenotypic probes, the cells were incubated with a sequencing primer (CACCTCATCCCACTCTTCAAAAGGACGAAACA CCG) at 1 μM concentration in 2X SSC with 10% formamide for 30 minutes at room temperature. Following this primer hybridization, the cells were washed three times with PR2 buffer (Nano kit PR2) and then incubated with incorporation mix (Nano kit reagent 1) for five minutes at 60°C. The incorporation mix was then removed over six serial dilutions with PR2 buffer. In order to decrease background fluorescence, the cells were washed with fresh PR2 buffer and incubated at 60°C for five minutes. The washing process was repeated five times before adding 200 ng/mL DAPI (Sigma-Aldrich #D9542-10MG) in 2X SSC and imaging.

### Fluorescence microscopy

Phenotypic and ISS images were acquired using a Nikon Ti-2 Eclipse inverted epifluorescence microscope with automated XYZ stage control, an Iris 9 sCMOS camera (Teledyne Photometrics), and hardware autofocus. All hardware was controlled using NIS-Elements AR, and a CELESTA light engine (Lumencor) was used for fluorescence illumination. Phenotypic images were acquired using a 20X 0.75 NA CFI Plan Apo Lambda objective (Nikon MRD00205) and the following Semrock filters for each phenotypic probe: Actin (phalloidin) emission ET530/30 nm, dichroic 495 nm. Mitochondria (TOMM20) emission 615/24 nm, dichroic 565 nm. Endoplasmic reticulum (Concanavalin A) emission 680/42 nm, dichroic 660 nm. Golgi and plasma membrane (WGA) emission 820/110 nm, dichroic 765 nm. Nucleus (DAPI) dual-band emission 408/473, dichroic. ISS cycles were imaged using a 10X 0.45 NA CFl Plan Apo Lambda objective (Nikon) with the following Semrock filters for each base: Miseq G excitation 543/4 nm, emission 575/30 nm, dichroic 555 nm. Miseq T emission 615/24 nm, dichroic 565 nm. Miseq A emission 680/42 nm, dichroic 660 nm. Miseq C emission 732/68 nm, dichroic 660. Laser power for all acquisitions was kept at 30%. Exposure times for ISS cycles were selected by balancing the average pixel intensities of *In Situ* Sequencing spots in each fluorescent channel.

### Next Generation Sequencing

Next generation sequencing was used for validation of plasmid libraries, cell libraries, and Cas9 activity in screening cell lines. For Cas9 activity assays and cell library validation, cell samples were lysed by resuspending cell pellets in lysis buffer (10 mM Tris pH 7.5, 1 mM CaCl^2^, 3 mM MgCl^2^, 1 mM EDTA, 1% Triton-X100, and 0.2 mg/mL Proteinase K), and heating for 10 minutes at 65°C followed by 15 minutes at 95°C. The target sequences in cell lysates were directly amplified without cell lysis purification according to the PCR reactions described below in Table 6. PCR amplification conditions were adapted from (Rana et al, Next-Generation Sequencing of Genome-Wide CRISPR Screens). Temperature conditions for PCR reactions followed initial denaturation at 95°C for 5 minutes, then denaturation at 95°C for 20 seconds, annealing at 55°C for 30 seconds, and extension at 72°C for 30 seconds. PCR 2 products were purified via gel extraction using the Qiaquick Gel Extraction Kit (Qiagen #28706X4) and prepared for sequencing as described in Illumina’s library denaturation and dilution manual. The PhiX Control library was spiked in the sequencing sample at 10% (v/v) (Illumina, cat. no. FC-110-3001).

**Table 6.**
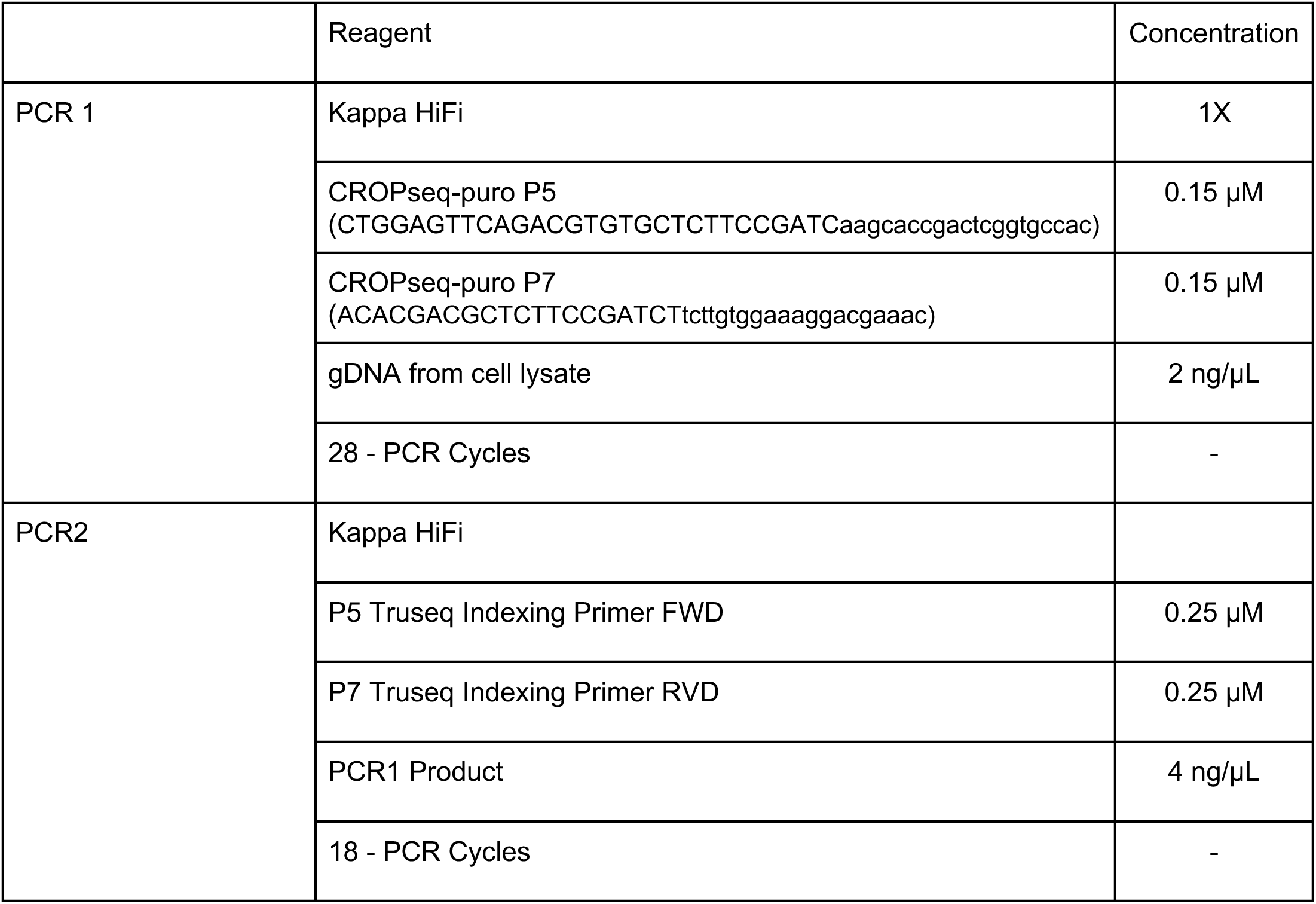
NGS library prep.

### Cell lines

The A549-TetR-Cas9 cell line was created by simultaneously transfecting A549 cells with *piggyBac* transposase and a *piggyBac* cargo plasmid containing TetR-inducible Cas9 (Addgene #134247), and selecting for 7 days with 500 ug/mL G418. Single cells were sorted into 96-well plates (Sony SH800) and expanded into colonies. An optimal clone was selected based on Cas9 activity, aiming for high and low activity in the presence and absence of doxycycline, respectively. Cas9 activity was evaluated using the fluorescence-based reporter pXPR011 (Addgene #59702), which expressed GFP and cognate sgRNA to assess GFP knockdown upon successful CRISPR activity. Fluorescence readouts of Cas9 activity were detected via FACS and Indel-Sequencing. The HeLa-TetR-Cas9 cell line was a gift from Iain Cheeseman; this cell line is a single cell clone selected for high Cas9 activity by transducing with the eGFP reporter mentioned above (pXPR011) and using FACS to read out efficiency of protein knockdown.

### Image processing

We used CellProfiler bioimage analysis software (version 4.1.3)^27^ to process the images using classical algorithms and Fiji (with openjdk-8)^68^ for image stitching^69^ and cropping. For the ISS images, we corrected for variations in background intensity, aligned channels within cycles, and performed channel compensation. For the phenotypic images, we corrected for variations in background intensity. We then stitched the ISS and CellPainting images independently into a full-well view and cropped them into corresponding pseudo-sites to account for the fact that they were imaged at different magnifications. Corrected, pseudo-site images from both ISS and phenotypic images entered our final analysis pipeline where they were aligned to each other, nuclei and cells were segmented using phenotypic images, ISS foci were identified and a barcode was called for each focus, and across the various channels captured, we measured various features of cells across several categories including fluorescence intensity, texture, granularity, density, and location (see http://cellprofiler-manual.s3.amazonaws.com/CellProfiler-4.1.3/index.html for more details). We obtained 3973 feature measurements from each of about 26.8 million (A549) and 46.4 million (HeLa) cells. We parallelized our image processing workflow using Distributed-CellProfiler^70^ and Distributed-FIJI^71^, triggered by Lambda Functions in Amazon Web Services. The actual CellProfiler pipelines used are available in the cellpainting-gallery (see Code and Data Availability) while continuously improved pipelines and Lambda Function scripts are available at https://github.com/broadinstitute/pooled-cell-painting-image-processing.

### Image-based profiling

We processed outputs of CellProfiler into image-based profiles using scripts available at https://github.com/broadinstitute/pooled-cell-painting-profiling-recipe. It is highly configurable beyond the configurations used for this report. The first step generates summaries of a variety of quality control metrics about the image acquisition, modified Cell Painting, and *in situ* sequencing. The second step uses pycytominer workflows to process the single cell features extracted using Cell Profiler. We median-aggregated the single cell profiles by guide for each plate independently. Next we defined the center and scale parameters as the median and median absolute deviation of feature values from non-targeting control perturbations, and then normalized the averaged profiles by subtracting the center value and scaling to the standard deviation, for each plate independently.

We further processed the per-plate guide level profiles to create the per-screen profiles we use in our analyses. We performed feature selection independently for each screen to eliminate noisy features and retain the most informative features by filtering out redundant features (all features that have Pearson correlation greater than 0.9 to a given feature), features with low variance, and features with missing values across all the plates as previously described^72^. Then we median-aggregated each experiment’s feature selected per-plate profiles to obtain a unique profile per guide for each experiment. For perturbation-level (gene-level) profiles, each experiment’s guide-level profiles were median-aggregated.

Each dataset is independently welded to the recipe, effectively versioning the recipe, using a Template, available at https://github.com/broadinstitute/pooled-cell-painting-profiling-template. Our A549 screen data with versioned recipe is available at https://github.com/broadinstitute/CP186-A549-WG. Our HeLa screens data with versioned recipe is available at https://github.com/broadinstitute/CP257-HeLa-WG. Code used for further profile processing is in this paper repository at https://github.com/broadinstitute/2022_PERISCOPE.

### Hit-calling, statistical analysis and distribution of hits

To determine the genes with significant signal above the noise (Hit-calling) we developed an algorithm to compare the distribution of values per feature for all the guides targeting the same gene to a set of non-targeting control guides using the Mann-Whitney U-test. The number of features significantly different from the non-targeting controls based on the statistical test (p-value 0.001) were added up to calculate profile score for each perturbation. Then to ensure that the perturbations called significant are truly not null we defined a control group called zero-TPM genes. Zero-TPM genes are the genes without significant expression in a given cell line (with zero transcript per million (TPM)) and were determined based on the RNA expression levels reported by the Broad Institute Dependency Map portal^31^. To obtain a false discovery rate (FDR) of 5%, perturbations with profile scores above 95% of zero-TPM genes were determined to have significant signal above the noise. Terms “whole-cell hits” and “compartment hits” were used to distinguish between perturbations with significant signal in overall profile features or perturbations with targeted signal in features from a specific cell compartment (based on one of the five fluorescent markers). For whole-cell hits all of the collected features were used in the hit-calling process explained above but for the compartment hits a subset of features from one cell compartment were used (including texture, intensity, correlation, radial distribution, and granularity measures from that compartment). It is important to note that a single perturbation can be a compartment hit, targeting simultaneously, two or rarely even three compartments but still not be a whole cell hit (see Extended Data Figure 3a,c,e).

### Distribution of significant features based on gene sets targeting each compartment

Pie charts showing the normalized fraction of number of features significantly different from the control categorized based on target compartments. The values are the average from multiple genes part of the highlighted gene groups.

### Comparison between pair-wise correlation of perturbations to other databases

To assess the ability of phenotypic profiles to recall known biological relationships, we calculated the correlation between profiles as a measure of similarity and used it to perform two global assessments. Considering the large number of features in each profile (1520 in A549,1597 in HeLa DMEM and 1709 in HeLa HPLM datasets) and to improve the signal to noise ratio, principal component analysis (PCA) was performed on the datasets to capture at least 70% of the variation. The resulting profiles were then used to calculate the Pearson correlation coefficient between all hit perturbation profiles (gene-level). First, annotated protein clusters were obtained from the 28.11.2022 CORUM4.0 database^29^. Clusters with at least 66% of the hit genes were identified using the gene symbols from both datasets (501 clusters in A549, 799 clusters in HeLa HPLM and 871 clusters in HeLa DMEM). Then all the correlations between each pair of genes in a cluster were calculated. The distribution of all the correlations between profiles within clusters versus the distribution of all the correlation between profiles from all hit genes were plotted in the figure. Second, we performed a similar analysis based on the protein link scores as predicted by the STRING database (v11.5, “9606.protein.links.v11.5.txt.gz”)^30^. To start, protein IDs from STRING were mapped to gene symbols using preferred_name extracted from the “9606.protein.info.v11.5.txt.gz” file. All the possible pairwise correlations between hit gene profile with a reported link score in the STRING database were calculated. Next, the correlations were binned into eight equally spaced bins and the distribution of the STRING link scores for each bin were plotted using seaborn.boxenplot^73^ in python.

### Comparison to Cancer Dependency Map Data

From DepMap data, we divided genes expressed in A549 cells into essential and nonessential categories based on Chronos gene effect scores^34^ using a threshold score of -0.5 for gene essentiality and plotted the distributions of essential and nonessential genes versus their morphological signal score.

### UMAP clustering of the hit perturbation profiles

To evaluate and demonstrate the ability of morphological profiles to uncover biologically relevant interactions and structures, UMAP (Uniform Manifold Approximation and Projection) algorithm was used to project the hit gene profiles into a 2-dimensional plane. PCA was performed on the datasets to capture at least 70% of the variation as described above before the application of the UMAP algorithm. The Python library UMAP was used to apply the UMAP algorithm using “cosine” for parameter “metric”. The details of the parameters used are available on the GitHub repository. Some of the resulting clusters were manually labeled to highlight some underlying interesting biology using gene ontology terms (biological processes and cellular components) as listed on the GSEA-MSigDB web portal (http://www.gsea-msigdb.org/gsea/msigdb/human/collections.jsp#C5).

### Hierarchical clustering of hit perturbation profiles and representative heatmaps

Correlations between morphological profiles is a powerful tool to extract biological insights from data sets. For example, similarity (or dissimilarity) contains information regarding functional clusters, protein structure, signaling pathways and their directionality. To this purpose, first, PCA was performed on the datasets to capture at least 70% of the variation as described above followed by the selection of a subset of perturbations associated with a functional gene-set as specified in each instance. Then the corr function from the pandas library in Python was used to calculate the pairwise Pearson correlation coefficient of the perturbation profiles for each dataset. The hierarchical clustering of the correlations and the plotting of the heatmaps was performed using the seaborn’s clustermap function in python. The ward variance minimization was used as the clustering algorithm (‘method’) based on the ’euclidean’ as the distance metric.

For combined heatmaps used in Figures 3h&i and Extended Data Figure 8b&c to compare DMEM and HPLM screens, the process explained above was performed on one screen as explained above (no heatmaps generated at this step). Then, the order of clustering was extracted from one screen and applied to the other screen to enable two types of comparisons: direct comparison between correlations from two screens and high level structural comparison in the clustered correlations. To effectively illustrate the output both sets of ordered correlations were merged into a single heatmap with the bottom left half representing one screen and the top right representing the other, using the seaborn.clustermap ^73^ function in python.

### Preranked GSEA analysis of perturbations based on morphological signal strength or similarity

To better understand the biological processes highlighted in each Hela screen and to compare the phenotypic downstream effects of the environment on cells, Preranked GSEA analysis was performed. The analysis was performed on the GSEA v.4.2.3 Mac software and the genes were ranked based on the morphological signal score using the “c5.go.bp.v2022.1.Hs.symbols.gmt [Gene ontology]” gene set database with 2000 permutations. The morphological signal score was calculated using this equation for each perturbation:

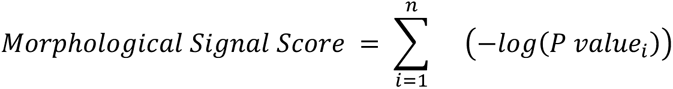

The p-values were calculated as described in the hit calling section and n refers to features significantly different from the non_targetting controls (p-value 0.001). The code used to calculate the morphological signal score as well as the list of perturbation scores for each dataset is available on the GitHub repository. The EnrichmentMap application based on the Cytoscape v3.9.1 software platform was used to visualize the enrichment maps (node cutoff Q-value 0.05).

Preranked GSEA analysis was performed to determine enrichment for biological terms based on morphological profile similarity to a query gene of interest. Genes were ranked based on cosine similarity to the profile of the query gene, then GO term enrichment was performed using the GSEApy package and the “GO_Cellular_Component_2021” database.

### Single Feature Screen Analysis

For each feature in the feature-selected dataset, genes were sorted by p-value (as generated during hit calling) and a Top 20+ list was created for each feature that contained all genes with a p-value less than or equal to that of the 20th gene. The Top 20+ list was assessed for GO term enrichment using the Python GOATOOLS library^74^ with the default Benjamini-Hochberg FDR correction. GO terms were considered enriched if they had a p-value <.05 after an additional Bonferonni correction. Compartment-specific gene lists were assayed for enrichment in the Top 20+ lists using a fisher exact test with a Benjamini-Hochberg FDR correction from the Python SciPy library^75^. Plots were made with Python library Matplotlib^76^. For exploration of Granularity features, guide-normalized but not feature-selected datasets were aggregated with pycytominer and plotted with Seaborn^73^. Gene lists were taken from the Metabolic Atlas^77^. Granularity features were visualized with Python SciPy and scikit-image^78^ libraries as implemented in CellProfiler.

### Atlas cell retrieval tool

Example single-cell image crops can be retrieved from any of the screens using a retrieval script included in our paper repository at https://github.com/broadinstitute/2022_PERISCOPE. Images are retrievable by gene name or sgRNA barcode sequence and example images can be chosen randomly or set to the most representative cells for that barcode as determined by closest k-means clustering using scikit-learn^79^. Individual channel crops are from corrected images on which the final analysis measurements are made. Mask crops are from segmentations generated during the analysis pipeline and are filled light gray to show the cell of interest and dark gray to show cells within the same crop assigned to the same perturbation.

### TMEM251 localization assay

HT1080 cells were transduced with lentiviral vectors expressing either TagBFP-tagged GALNT2 (Golgi) or mRFP1-tagged TMEM192 (Lysosome), and selected with antibiotics. Cells with stable integration were fixed with 4% formaldehyde (15 minutes at 4°C), permeabilized with 20 ug/mL digitonin (30 minutes at room temperature), blocked with 1% BSA (30 minutes at room temperature), and incubated with primary antibody against TMEM251 (HPA048559, Sigma-Aldrich; 1:200 overnight at 4°C) followed by Alexa Fluor 488-conjugated secondary antibody (1:1000, 2 hours at room temperature). Samples were imaged on the Phenix imager (Perkin-Elmer) with a 63x objective in confocal mode.

### WGA / LAMP1 co-staining and quantification of lysosomal glycan accumulation followed by CRISPRi perturbations

HT1080 CRISPRi cells were transduced with sgRNA-expressing lentiviral vectors and selected with antibiotics. For dual-target samples, cells were transduced simultaneously with two vectors and co-selected with two antibiotics. 8 days after sgRNA transduction and 2 days after final re-plating, cells were fixed, permeabilized, blocked, and stained as above, using primary antibody against LAMP1 (ab25630, Abcam; 1:50) and Alexa Fluor 647-conjugated secondary antibody. Alexa Fluor 555-conjugated WGA at 1.5 ug/mL and Hoechst 33342 at 5 ug/mL were included during secondary antibody incubation. Samples were imaged on the Phenix imager (Perkin-Elmer) with a 63x objective in confocal mode.

Image analysis was performed using the Harmony software (Perkin-Elmer), where images were flat-field corrected and regions corresponding to the nucleus, cytoplasm, and lysosome were identified. WGA signals that colocalized with the lysosomes were quantified by the median fluorescence intensity (MFI) for each cell. Each biological replicate (two per condition) was represented by the upper quartile of the per-cell MFIs from all segmented cells.

### Lysosomal pH measurement

HT1080 CRISPRi cells stably expressing rat Lamp1 tagged with mScarlet (on the lumen side) were transduced with sgRNA-expressing lentiviral vectors and selected with antibiotics. Cells were imaged live, in an environmental control chamber (OKO) at 37°C and 5% CO^2^, 8 days after sgRNA transduction and 1 day after re-plating into Imaging Media on an 8-well chambered cover glass (C8-1.5H-N, Cellvis). Imaging was performed on an SP8 scanning microscope (Leica) in FLIM mode using a 100x objective. Samples were excited by a white light laser at 561 nm and 40MHz, and emission collected between 590-700 nm. Imaging Media consisted of FluoroBrite DMEM (A1896701, Life Technologies) + 10% FBS + 1% GlutaMax (35050061, Gibco).

Image analysis was performed using in-house scripts, which identified lysosomal regions and the mean arrival time (lifetime) of photons in each pixel. The median lifetime from all lysosomal pixels in each field-of-view (consisting of 1-2 cells each, with ≥15 fields per condition) was computed and represented as one data point per field-of-view. After the initial imaging, 100 nM Bafilomycin A1 were added to the Non-targeting sample for a positive control, which was reimaged 5 hours after the treatment.

### Lysosomal hydrolase activity assay

HT1080 CRISPRi cells were transduced with sgRNA-expressing lentiviral vectors and selected with antibiotics. 9 days after sgRNA transduction and 1 day after final re-plating, cells were assayed for their lysosomal hydrolase activity by incubating with 0.2 ug/mL Hoechst 33342 and either 200 uM PFB-FDGlu (for glucosylceramidase; P11947, Invitrogen) or 33 uM C^12^FDG (for beta galactosidase; I2904, Invitrogen) in Imaging Media for 1 hour at 37°C, before imaging on the Phenix imager (Perkin-Elmer) with a 63x objective in confocal mode.

Image analysis was performed using the Harmony software (Perkin-Elmer), where flat-field corrected images were segmented for nucleus and cytoplasm. Total fluorescence intensity for each cell was extracted, and each biological replicate (two per condition) was represented by the median of the per-cell fluorescence (MFI) from all segmented cells, relative to the Non-targeting controls, as log10 fold-change.

### Code and Data availability

All code and data are publicly available. Phenotyping and in situ sequencing images and image-based profiles are available at the Cell Painting Gallery on the Registry of Open Data on AWS (https://registry.opendata.aws/cellpainting-gallery/) under accession number cpg0021-periscope. Instructions for retrieving images and profiles are available within the Cell Painting Gallery documentation at https://github.com/broadinstitute/cellpainting-gallery. The exact CellProfiler pipelines used in the screen are available in the Cell Painting Gallery while continuously improved image analysis pipelines and AWS Lambda scripts used to trigger them are available at https://github.com/broadinstitute/pooled-cell-painting-image-processing. Image based profiling data is welded to individual datasets using a template available at https://github.com/broadinstitute/pooled-cell-painting-profiling-template. It is processed with a recipe available at https://github.com/broadinstitute/pooled-cell-painting-profiling-recipe. The recipe outputs for the datasets that we report here are available at https://github.com/broadinstitute/CP186-A549-WG and https://github.com/broadinstitute/CP257-HeLa-WG.

The recipe outputs were further processed to generate the profiles analyzed in this paper. Code for the final processing and the creation of all figures in this paper are available at https://github.com/broadinstitute/2022_PERISCOPE.

## Acknowledgements

We thank all members of the Cimini, Carpenter-Singh, Blainey, and Neal labs for helpful discussions and feedback. We thank Iain Cheeseman for the HeLa-TetR-Cas9 cell line. The HeLa cell line was used in this research. Henrietta Lacks, and the HeLa cell line that was established from her tumor cells without her knowledge or consent in 1951, has made significant contributions to scientific progress and advances in human health. We are grateful to Lacks, now deceased, and to the Lacks family for their contributions to biomedical research. This work was supported by Calico Life Sciences LLC., the Novo Nordisk Foundation (NNF21SA0072102, M.C. & J.T.N.), and NIH grant no. 1DP2GM146252 (J.T.N.).

## Contributions

Conceptualization: J.T.N., P.C.B., A.S., A.E.C., S.S., B.A.C., C.H.J.. Formal analysis: M.R., S.S., B.A.C., E.W., G.P.W., J.Y.. Funding acquisition: J.T.N., C.H.J.. Investigation: C.D., J.B., S.L.K., M.E.L., A.S., J.Y., M.H.. Methodology: J.T.N., J.B., M.R., P.C.B., A.S., A.E.C., S.S., B.A.C., E.W., G.P.W., J.Y.. Software: M.R., A.E.C., S.S., B.A.C., E.W., G.P.W., M.H.. Supervision: J.T.N., P.C.B., A.S., A.E.C., S.S., B.A.C., G.P.W., C.H.J., M.C..Validation: M.R., M.E.L., J.Y.. Visualization: M.E.L., M.R., E.W., J.Y., M.H.. Writing - original draft: J.T.N., M.E.L., M.R., E.W., C.H.J., J.Y.. Writing - review & editing: J.T.N., C.D., J.B., M.E.L., M.R., P.C.B., A.S., A.E.C., S.S., B.A.C., E.W., G.P.W., C.H.J., J.Y.., M.C.,J.P-S., T.B..

## Extended Data

**Extended Data Figure 1.**
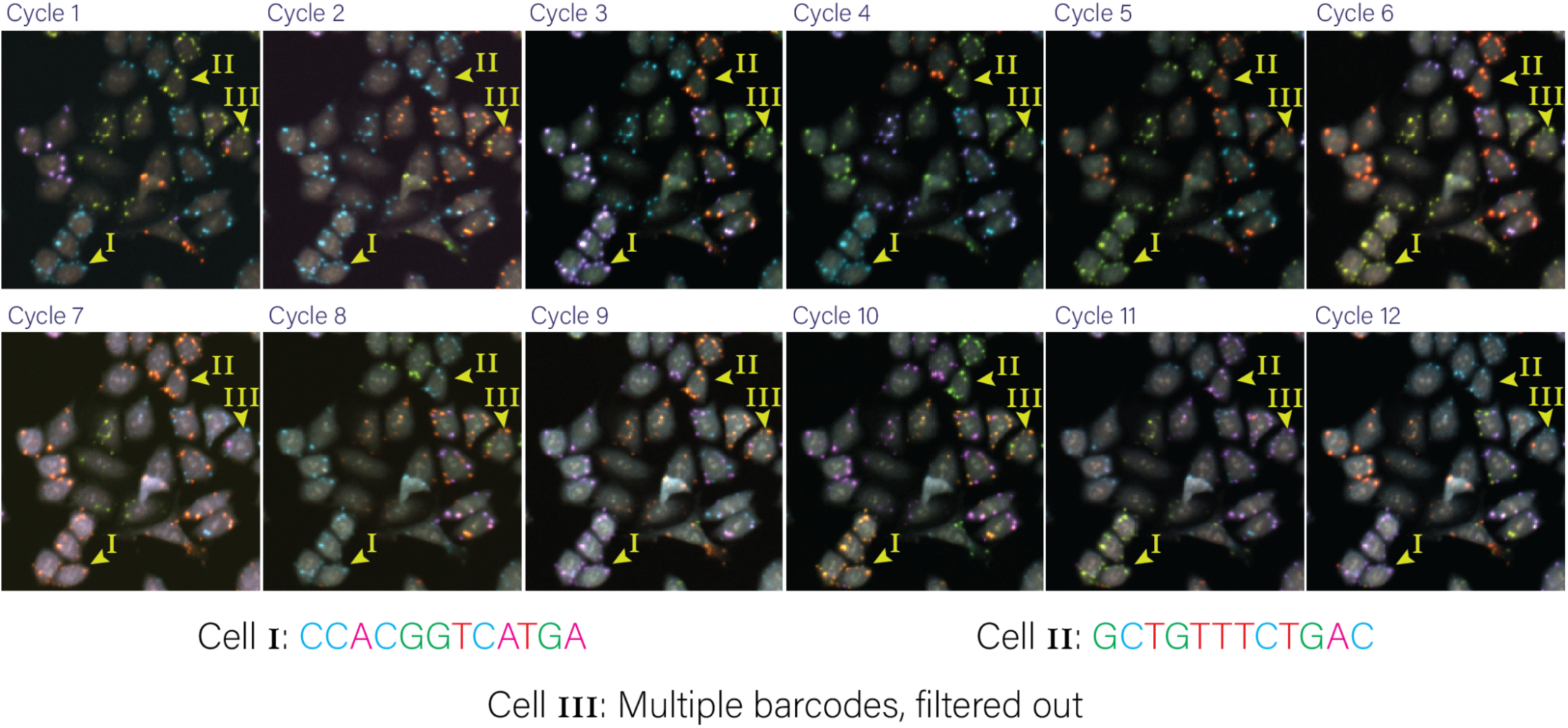
Example barcode calling based on twelve in-situ cycles. An example of a group of cells tracked over the twelve cycles of in-situ sequencing to call barcodes. Cells 1 and 2 highlight how the signal from fluorescent nucleotides are translated into a barcode read over twelve cycles.

**Extended Data Figure 2.**
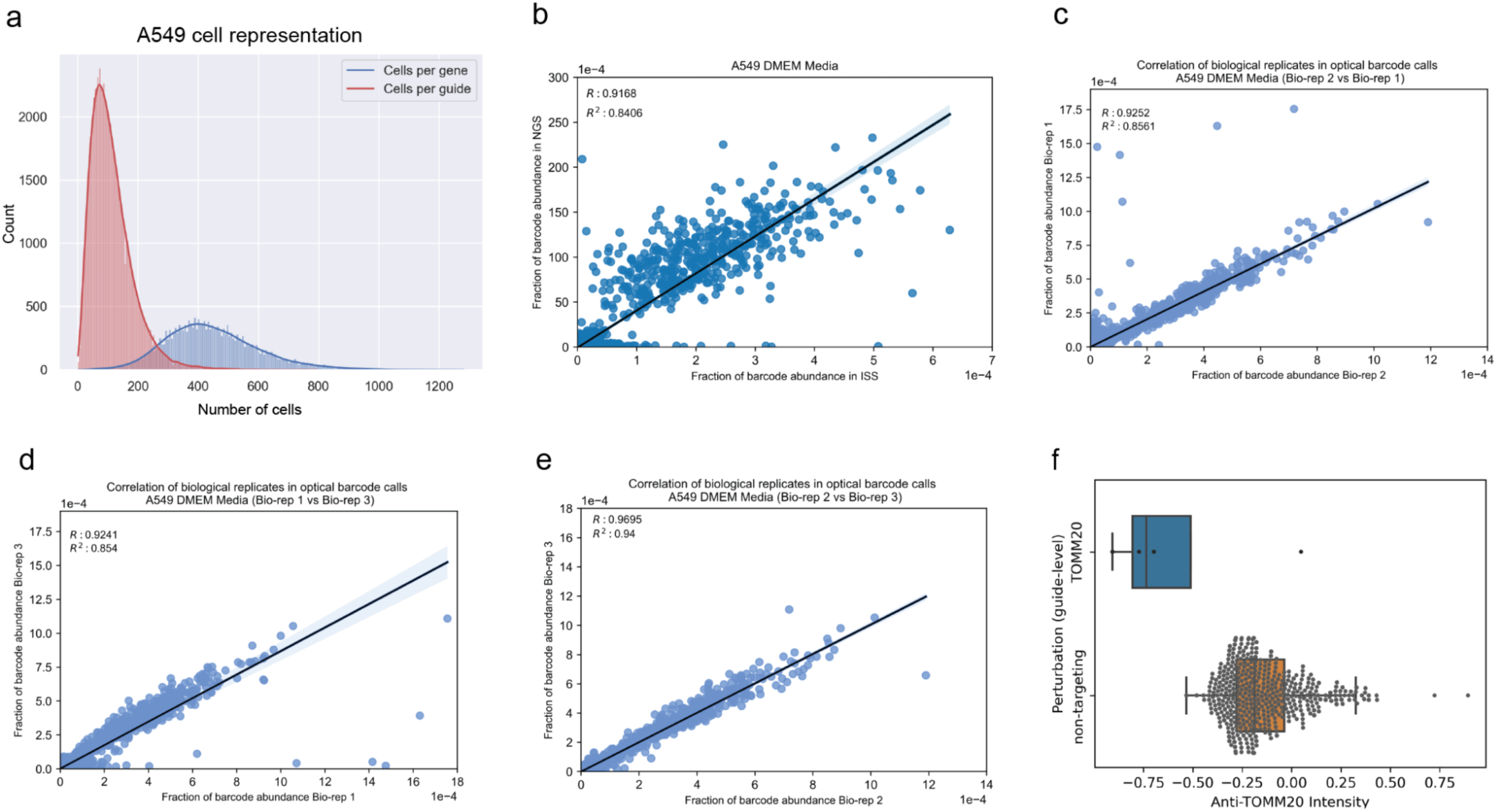
Technical summary of the A549 whole genome screen. (a) The distribution for the number of cells per gene or per guide present in the A549 dataset. (b) Comparison of the relative abundance of barcodes as quantified by NGS or in situ sequencing (R^2^ = 0.84). (c-e) Comparison of the relative abundance of barcodes as quantified by in situ sequencing among 3 different bioreplicates representing individual viral transductions (R_12_^2^ = 0.85, R_13_^2^ = 0.85, R_2_^32^ = 0.94). (f) The distribution of normalized mean mitochondria channel intensity per cell for guides targeting the TOMM20 gene and the non-targeting control guides.

**Extended Data Figure 3.**
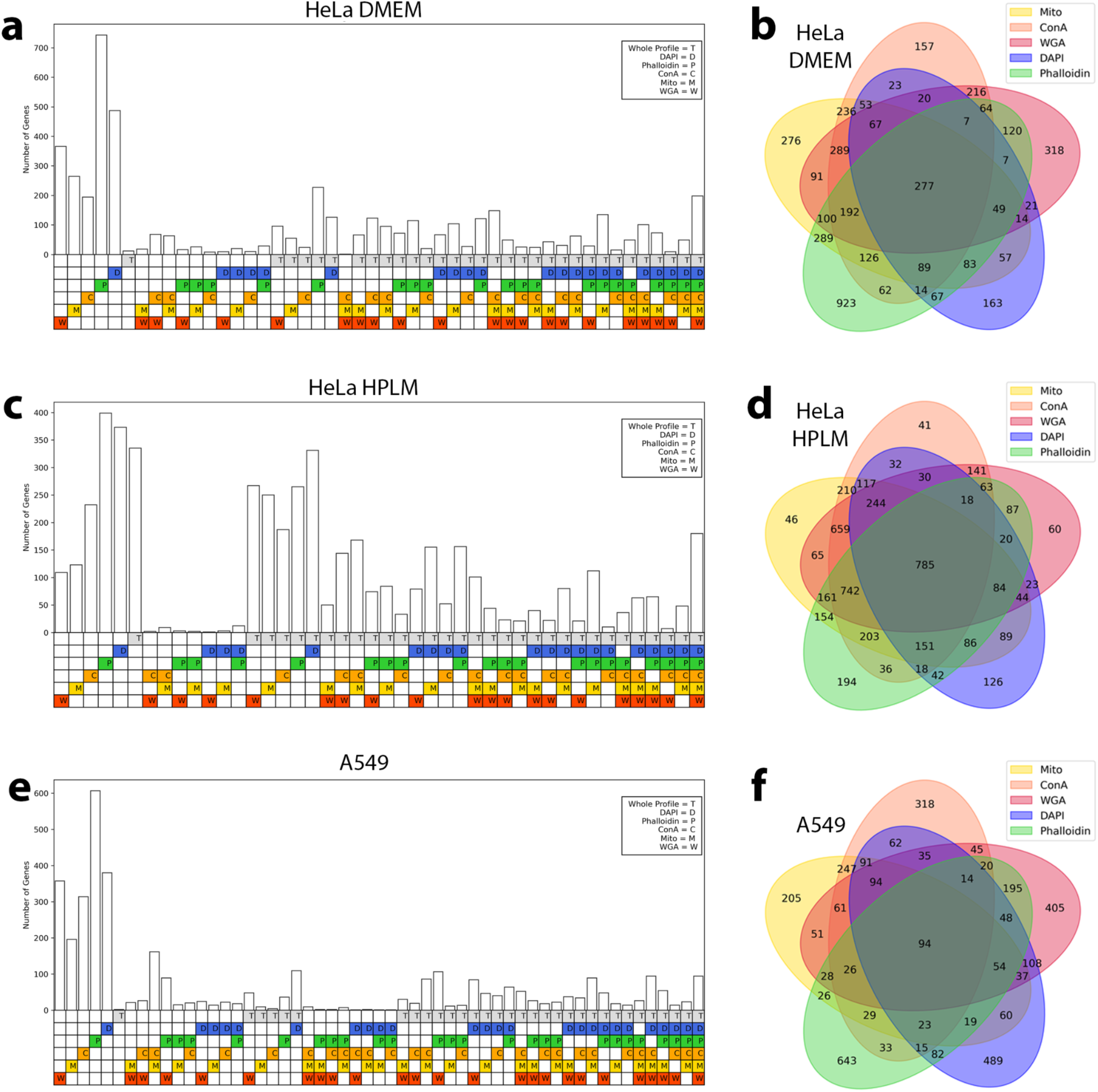
Hit genes can be called in multiple channel combinations. Genes called as hits in the HeLa DMEM (a-b), HeLa HPLM (c-d), and A549 (e-f) screens can be called as hits because of significant perturbation to their whole profile, any individual screen channel, or any combination thereof. Specific combinations without any hit genes are omitted from the bar plots (a,c,e) and whole profile hit information is omitted from the Venn diagrams (b,d,f) for clarity.

**Extended Data Figure 4.**
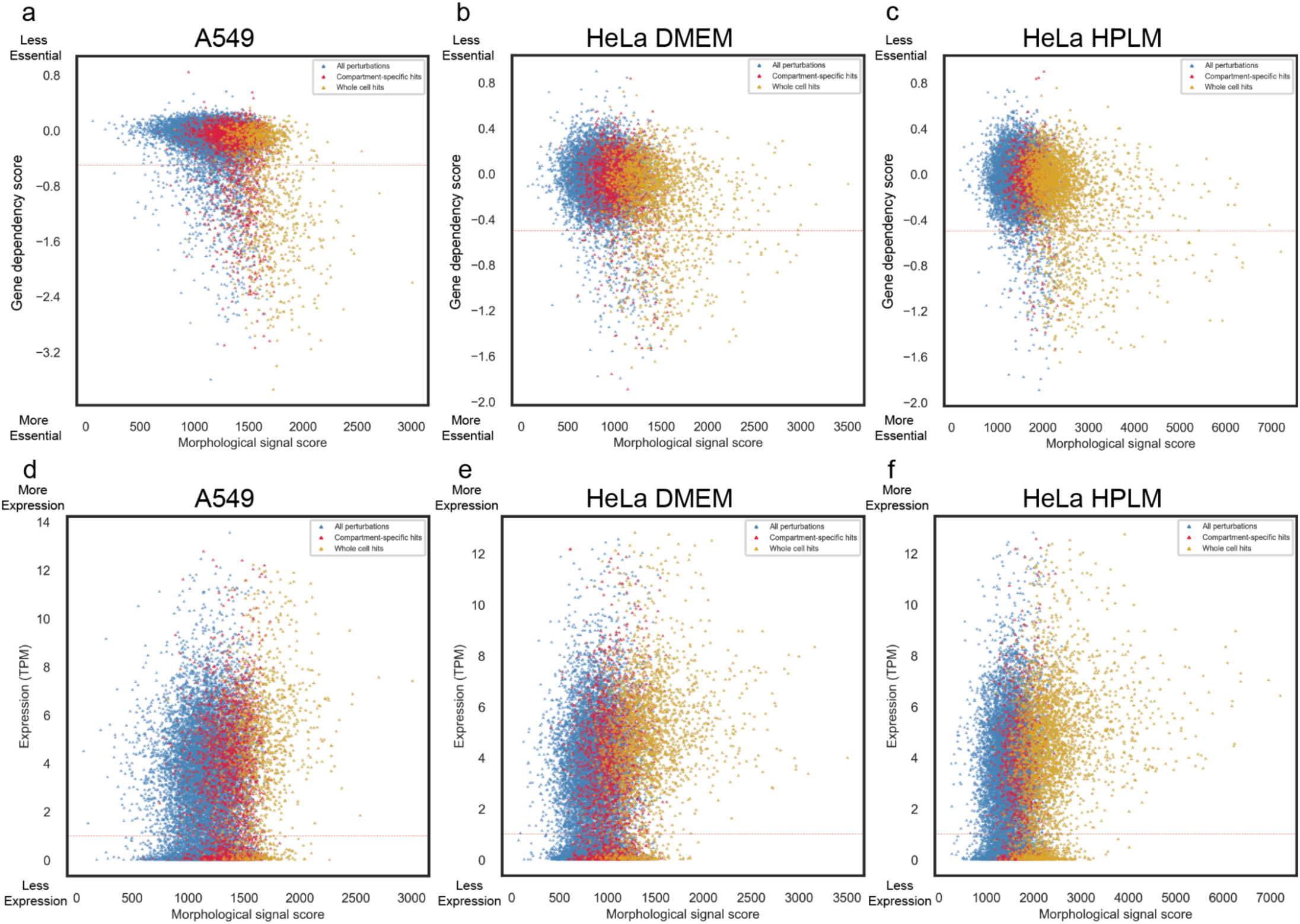
Morphological signal score is not well correlated with gene dependency or baseline gene expression. Comparison of the distribution of morphological signal scores and gene dependency scores (DepMap gene effect for the A549 cells and genetic dependencies estimated using the DEMETER2 for HeLa cells, the dashed red line at -0.5 threshold to highlight likely essential genes) for the A549 (a), the HeLa DMEM (b) or HeLa HPLM dataset (c). Comparison of the distribution of morphological signal scores and gene expression TPM values (DepMap dataset, values are inferred from RNA-seq data using the RSEM tool) for the A549 (d), the HeLa DMEM (e) or HeLa HPLM dataset (f).

**Extended Data Figure 5.**
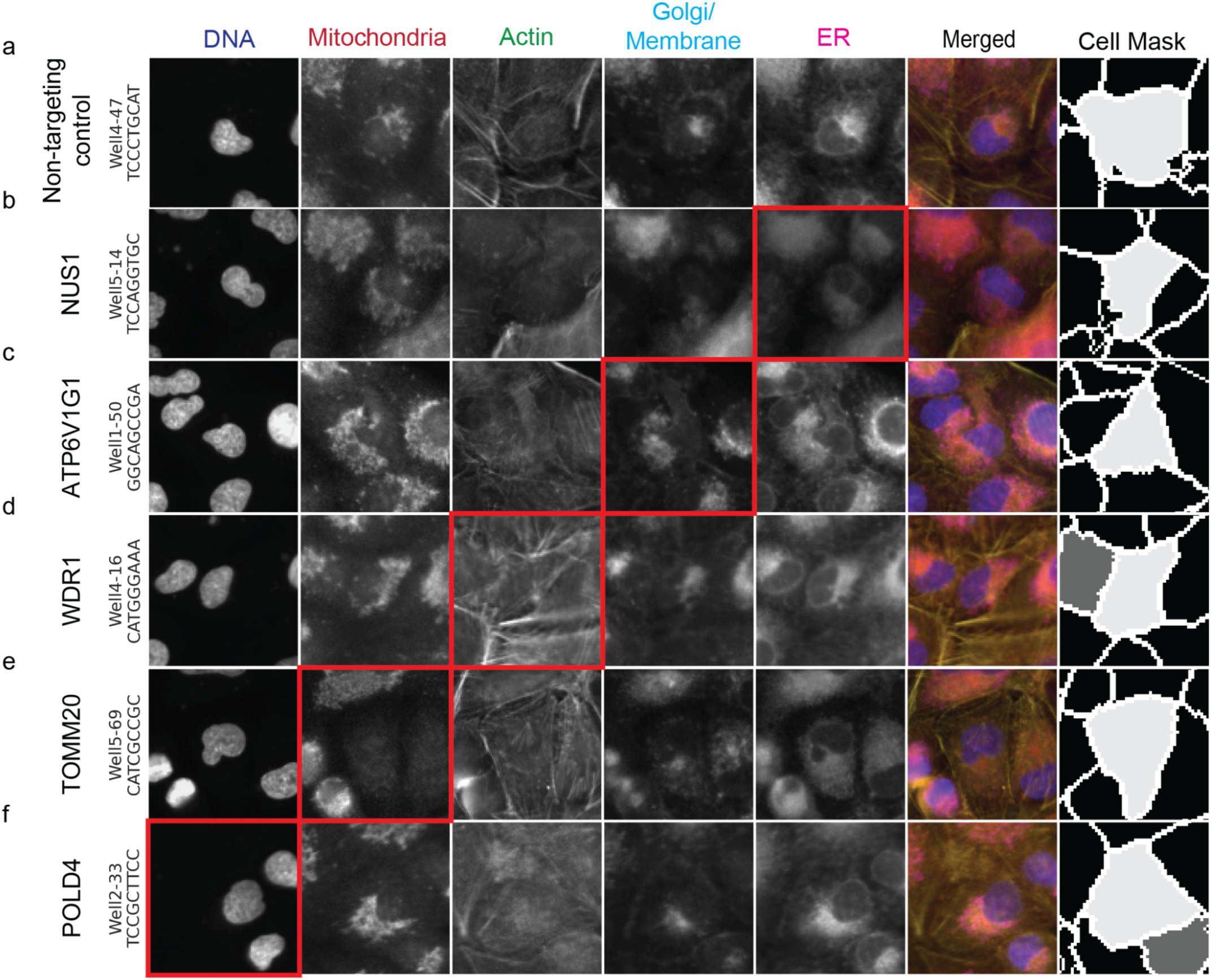
Examples of single cell images with morphological phenotypes labeled. Single cell images of specific perturbations retrieved from the A549 dataset representing five separate compartments and the segmentation shown as the cell Mask. (a) Single cell expressing non-targeting control guide RNA. Panels (b) to (f) represent images of single cells carrying guide RNA targeting genes with significant signal in specific cell compartment highlighted by the red box (Genes were selected based on the number of significant features targeting the specified compartment from gene sets highlighted in the Figure 2c).

**Extended Data Figure 6.**
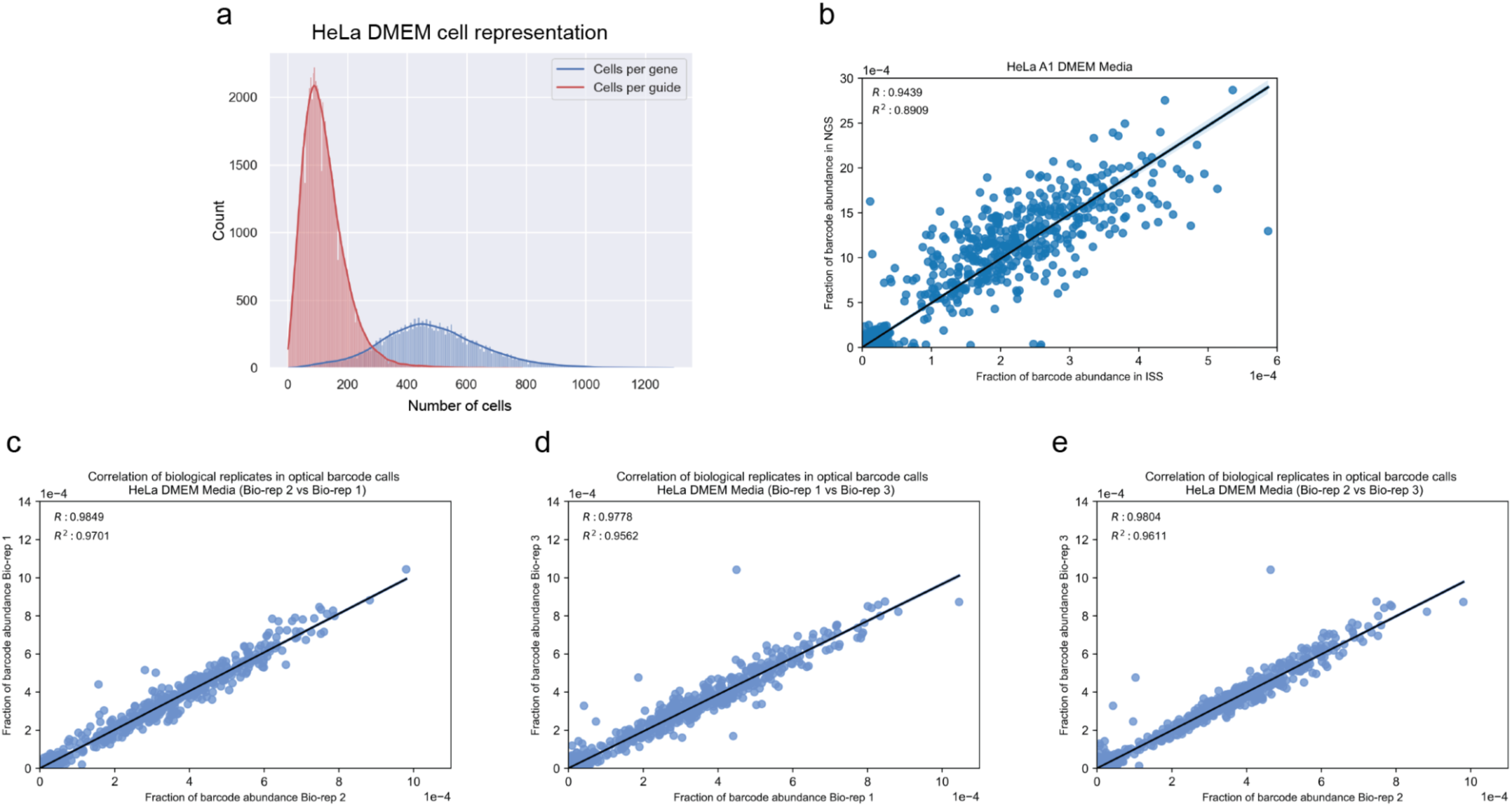
Technical summary of the DMEM HeLa whole genome screen. (a) The distribution for the number of cells per gene or per guide present in the DMEM HeLa dataset. (b) Comparison of the relative abundance of barcodes as quantified by NGS or in situ sequencing (R^2^ = 0.89). (c-e) Comparison of the relative abundance of barcodes as quantified by in situ sequencing among 3 different bioreplicates representing individual viral transductions (R_1_^22^ = 0.97, R_1_^32^ = 0.95, R_2_^32^ = 0.96).

**Extended Data Figure 7.**
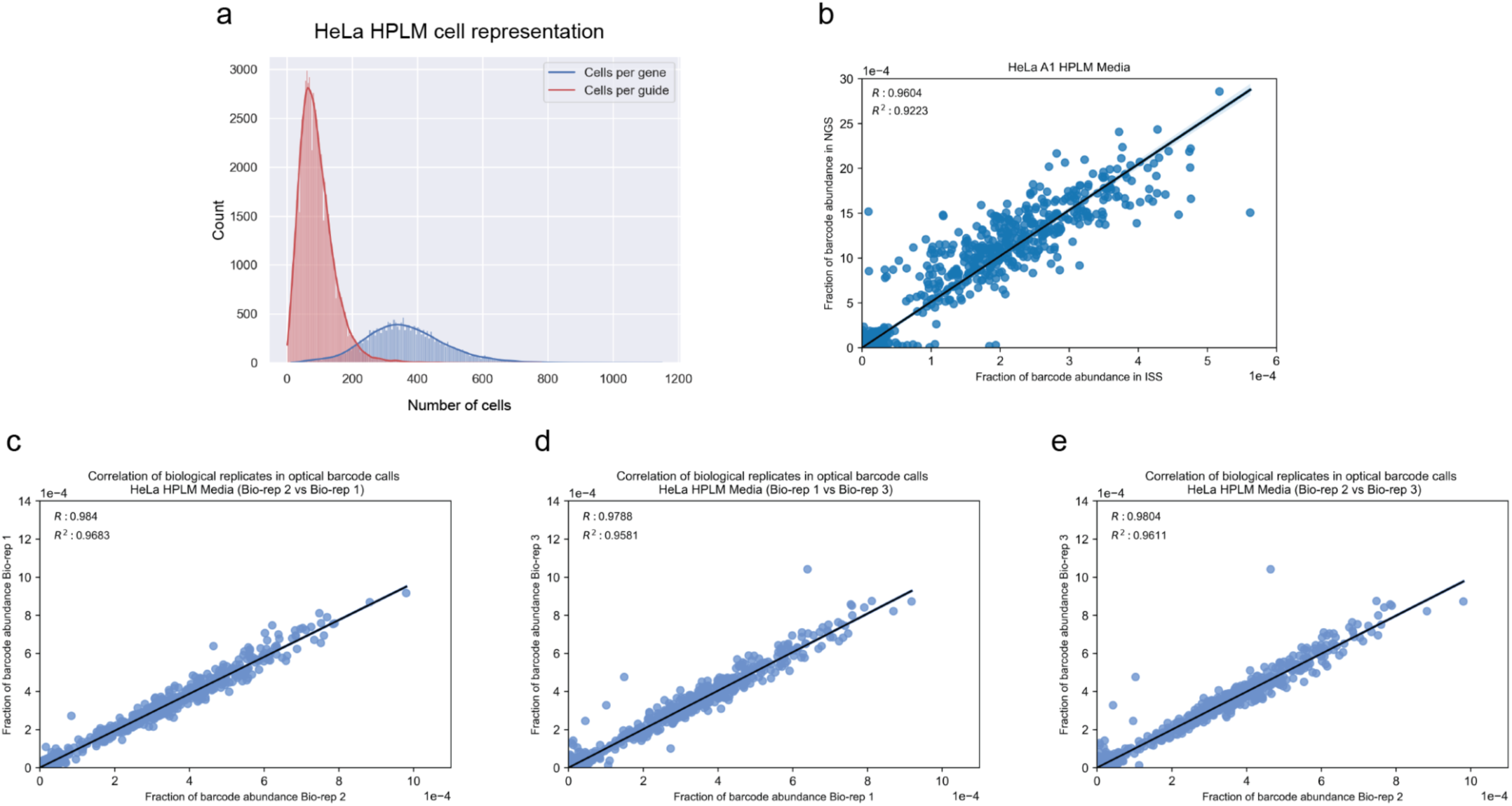
Technical summary of the HPLM HeLa whole genome screen. (a) The distribution for the number of cells per gene or per guide present in the HPLM HeLa dataset. (b) Comparison of the relative abundance of barcodes as quantified by NGS or in situ sequencing (R^2^ = 0.92). (c-e) Comparison of the relative abundance of barcodes as quantified by in situ sequencing among 3 different bioreplicates representing individual viral transductions (R_1_^22^ = 0.96, R_1_^32^ = 0.95, R_2_^32^ = 0.96).

**Extended Data Figure 8.**
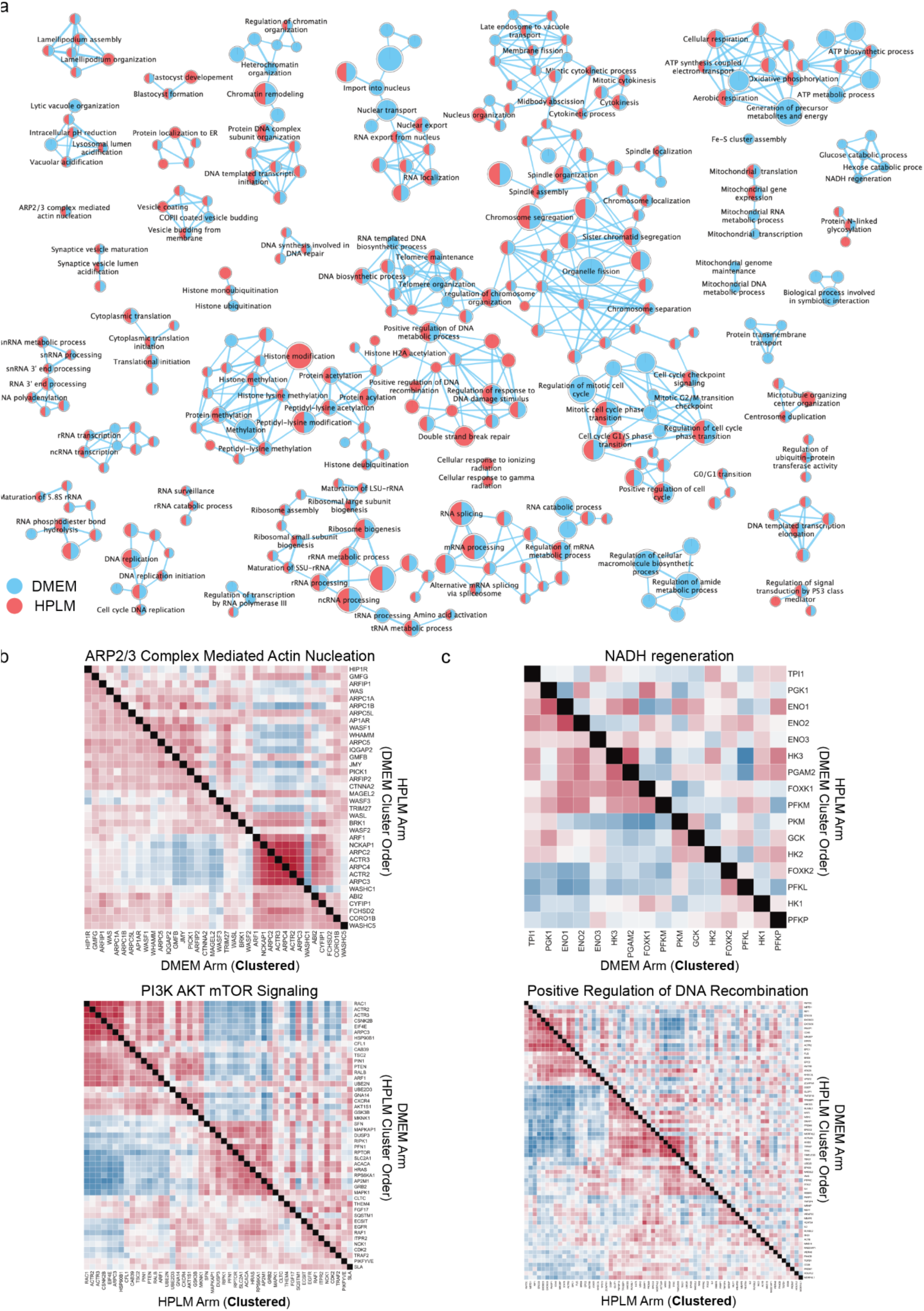
PERISCOPE identifies media-specific perturbation signatures. (a) The enrichment map for biological processes based on the GSEA analysis of the profile signal strength from the DMEM and HPLM HeLa screens. The preranked GSEA analysis was performed using a list of all genes ordered based on the calculated signal strength as described in methods. The Gene Ontology Biological Processes (GOBP) gene set was employed for the enrichment analysis. Some of the labels and single/double nodes are not shown here for clarity (Full map available in the supplemental figures). (b, c) Heatmaps representing Pearson’s correlation between gene profiles after hierarchical clustering using Ward’s method. Gene complexes/processes were enriched in both HeLa DMEM and HPLM datasets (b) or one dataset (c, the enriched screen is identified by **Clustered**)based on the preranked GSEA analysis. The heatmaps are a combination of Pearson’s correlation from both screens and clustered based on the data from a single screen as described in the cartoon in Fig 3 (g).

**Extended Data Figure 9.**
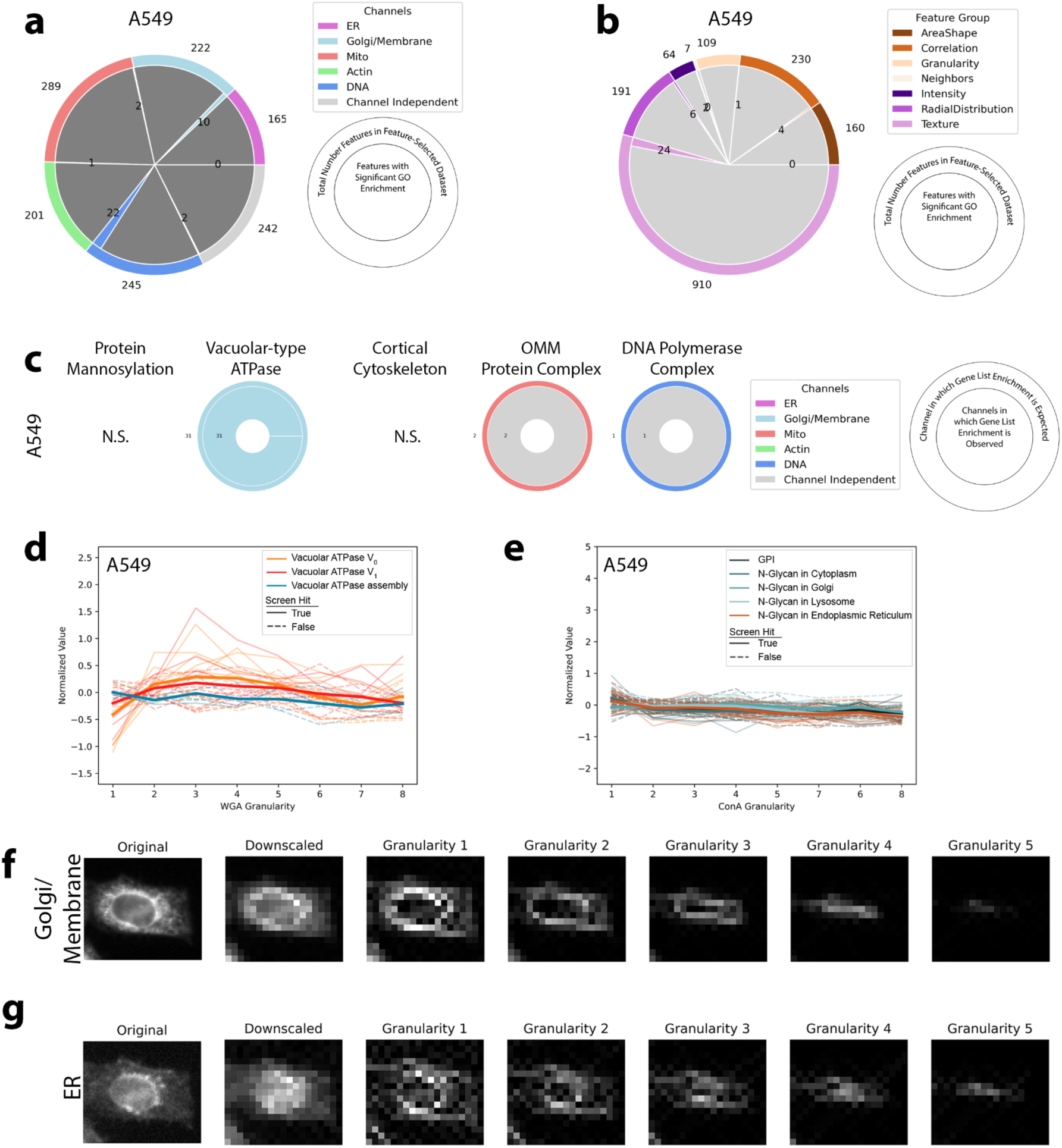
Identifying biological pathways using individual subcellular image features in A549 dataset. The A549 dataset shows minimal GO enrichment in individual features making distribution of enrichment across channels (a) and feature categories (b) difficult to interpret. Outer ring is the total number of features in our feature-selected dataset and inner ring is the number of features that showed GO enrichment for a and b. (c) Vacuolar ATPase protein products are expected to function specifically in the WGA channel and vATPase genes are specifically enriched in hit lists for features in those compartments. N.S. indicates no enrichment in that gene list. Outer ring indicates the channel in which enrichment is expected. Inner ring is the breakdown of actual channels that show enrichment for the gene group. (d) Disruption of the Vacuolar ATPase (either V^0^ or V^1^ subunit) but not genes involved in its assembly causes a decrease in Golgi/Membrane signal in small structures as seen specifically with screen feature WGA_Granularity_1 but not larger granularities. Each trace is a single gene; those genes that are not hits in the screen are dashed. Bold lines are the mean of all genes in the group. (e) Specific signal in granularity features is not observed for a loss of function in genes involved in N-Glycan synthesis in the Endoplasmic Reticulum. Visualization of the signal measured at each granularity is shown for Golgi/Membrane (f) and Endoplasmic Reticulum (g).

**Extended Data Figure 10.**
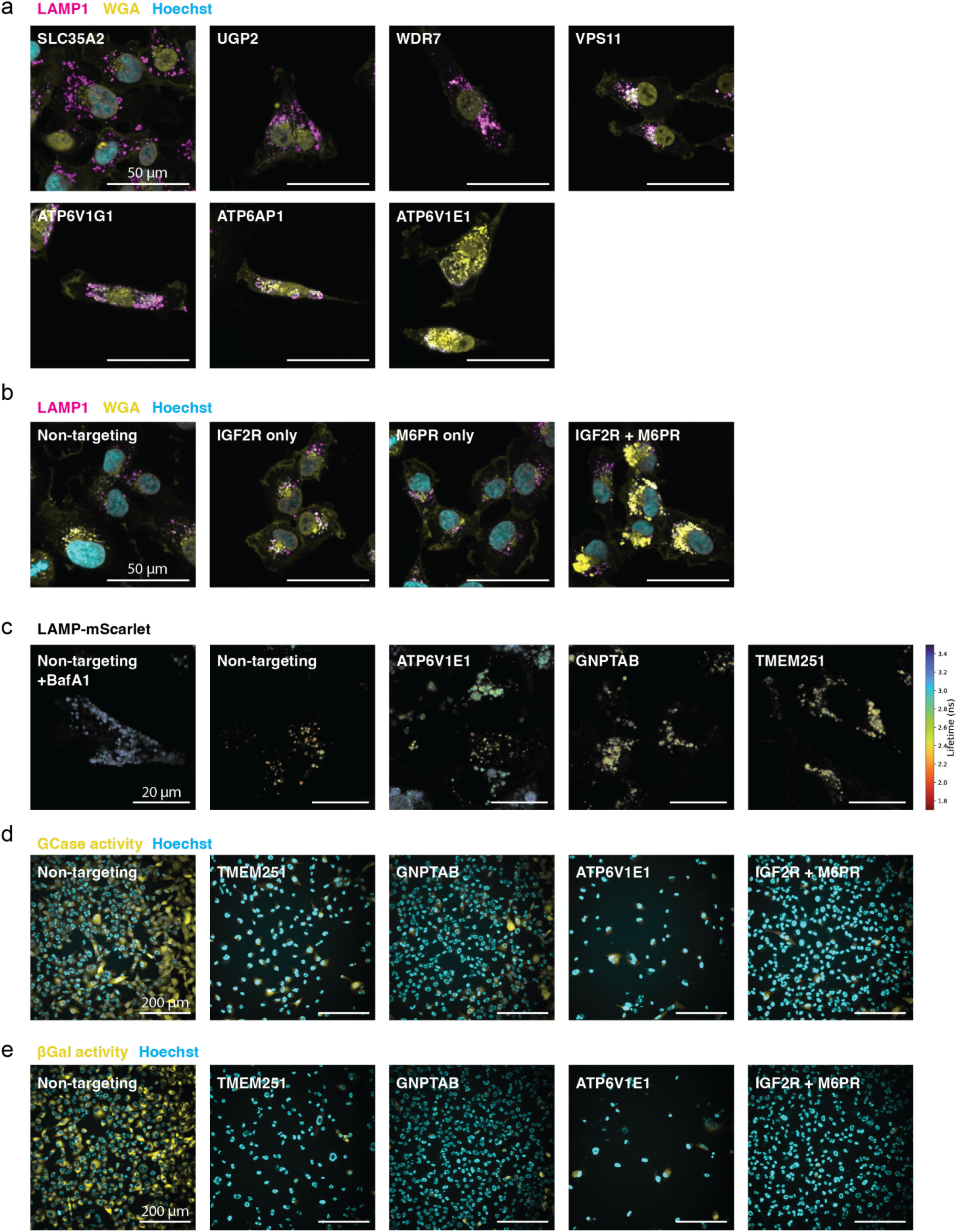
Phenotypic consequences of lysosomal trafficking perturbations (sample images contributing to Figure 6) (a) Confocal images of cells co-stained with WGA and LAMP1 antibody, as in Figure 6d, with KD of the remaining genes highlighted in Figure 6b as indicated. Quantified data were shown in Figure 6e. (b) Confocal images of cells co-stained with WGA and LAMP1 antibody, with single or dual gene KD as indicated. Quantified data were shown in Figure 6f. (c) Color overlays of mScarlet-Lamp1 cells with KD of genes indicated. Image intensity represents photon count per pixel, whereas hue encodes median lifetime per pixel. Quantified data were shown in Figure 6g. (d-e) Confocal images of live cells stained with KD of genes indicated and incubated with fluorogenic substrates of glucosylceramidase (d) and beta galactosidase (e), respectively. Quantified data were shown in Figure 6h-i.

